# Structural insights into mitotic-centrosome assembly

**DOI:** 10.1101/2025.10.03.680336

**Authors:** Nada Mohamad, Siu-Shing Wong, Anupa Majumdar, Alan Wainman, Ingelise Holland-Kaye, Lars Hubatsch, Zsofia Novak, Andrey Pozniakovsky, Martine Ruer-Gruss, Andreas F.M. Haensele, Anna Caballe, Steven Johnson, Susan M. Lea, Anthony Hyman, Jordan W. Raff

**Affiliations:** Sir William Dunn School of Pathology, University of Oxford, Oxford OX1 3RE, UK; The Max Planck Institute of Molecular Cell Biology and Genetics, Dresden, Germany; St. Jude Children’s Research Hospital

## Abstract

Centrosomal material assembles rapidly in mitosis. In *Drosophila*, the coiled-coil protein Cnn forms a scaffold that recruits PCM clients; in *C.elegans*, SPD-5 plays an analogous role. Here we show that full-length Cnn and SPD-5 can both form spherical condensates *in vitro*, but that the interactions driving their assembly into scaffolds inside cells appear to diverge. We show that the Cnn PReM adopts a helical hairpin fold that autoinhibits CM2 binding but that phosphorylation appears to increase hairpin breathing to permit CM2 engagement and robust scaffold assembly. Phospho-blocking mutations prevent PReM–CM2 interactions and scaffold formation, whereas phospho-mimetic substitutions partially restore function. The human homologue CDK5RAP2 contains a CM2 domain that can partially substitute for fly CM2 *in vivo* and we identify a candidate CDK5RAP2 PReM region that forms macromolecular networks with human CM2 *in vitro*. By contrast, the putative PReM and CM2 regions of SPD-5 cannot substitute for their equivalent fly domains and they do not interact detectably, suggesting a distinct assembly mechanism in worms despite conserved PLK1-dependent control of PCM growth.

## Introduction

Centrosomes have many important functions, and their dysfunction has been linked to a plethora of human pathologies, including cancer and microcephaly. Centrosomes are formed when a central pair of centrioles recruit a matrix of pericentriolar material (PCM) around themselves (Conduit et al., 2015; Bornens, 2021; Vasquez-Limeta and Loncarek, 2021; Gomes Pereira et al., 2021). During cell division, the amount of PCM at the centrosomes increases dramatically in a process termed centrosome maturation (Palazzo et al., 2000). This allows centrosomes to serve as dominant sites of MT nucleation during cell division, forming the two poles of the mitotic spindle that segregates the cell’s genetic material (Meraldi, 2016).

The mitotic PCM probably comprises several hundred proteins (Alves-Cruzeiro et al., 2014; Huang et al., 2015), but despite this molecular complexity, it can assemble and disassemble very rapidly as cells prepare to enter and exit mitosis, respectively (Enos et al., 2018; Magescas et al., 2019; Mittasch et al., 2020; Wong et al., 2024). This has prompted much debate about the mitotic-PCM’s biophysical nature, as it must be strong enough to resist the forces exerted by the MTs it organises and also provide an environment in which hundreds of proteins can be concentrated and potentially interact with one another. In particular, it remains unresolved as to whether liquid-liquid phase separation (LLPS) may be important for mitotic centrosome assembly (Rale et al., 2018; Woodruff, 2018; Raff, 2019; Lee et al., 2021; Woodruff, 2021), as has been suggested for several other cellular condensates (Shin and Brangwynne, 2017; Alberti and Hyman, 2021).

Although flies and worms are separated by ∼800-1000 million years of evolution (Erwin et al., 2011), a similar set of molecules are required for mitotic centrosome assembly. In both species the centriole and PCM protein Spd-2/SPD-2 (fly/worm nomenclature) (O’Connell et al., 2000; Kemp et al., 2004; Dix and Raff, 2007; Giansanti et al., 2008) recruits the mitotic kinase Polo/PLK-1 (Decker et al., 2011; Cabral et al., 2019; Alvarez Rodrigo et al., 2019; Wong et al., 2022), which then phosphorylates either Cnn (flies) or SPD-5 (worms) to stimulate the assembly of a macromolecular “scaffold” that ultimately recruits most, if not all, of the other PCM “client” proteins to the mitotic centrosome (Decker et al., 2011; Woodruff et al., 2015; Cabral et al., 2019; Ohta et al., 2021; Conduit et al., 2014a; b). Fly and worm Spd-2/SPD-2 and Polo/PLK-1 are clear homologues, but Cnn and SPD-5 share little sequence homology—although they are both predicted to be large coiled-coil-rich proteins. Thus, it remains unclear whether these two, largely unrelated, molecules form mitotic-PCM scaffolds that assemble and function in a similar manner.

The internal molecular interactions that drive Cnn and SPD-5 scaffold assembly are beginning to be elucidated (Feng et al., 2017; Nakajo et al., 2022; Rios et al., 2024). Cnn has two conserved motifs (Zhang and Megraw, 2007): CM1, which recruits and regulates the γ-tubulin ring complex (γ-TuRC) (Tovey et al., 2021; Serna et al., 2024; Dendooven et al., 2024; Gao et al., 2025), and CM2, which has been proposed to help target this family of proteins to centrosomes (Wang et al., 2010; Citron et al., 2018) (Figure 1A). In flies the CM2 domain interacts with an internal **P**hospho-**Re**gulated **M**ultimerization (PReM) domain that contains several potential Polo/PLK1 phosphorylation sites and is centred around a **L**eucine **Z**ipper (LZ) whose crystal structure when bound to CM2 (in a tetrameric LZ_dimer_::CM2_dimer_ complex) has been solved (Feng et al., 2017). When the full PReM and CM2 domains are mixed *in vitro*, they form large micron-scale assemblies and point mutations that perturb the LZ::CM2 tetramer perturb PReM::CM2 scaffold assembly *in vitro* and Cnn scaffold assembly *in vivo*. CDK5RAP2, the human homologue of Cnn, has both CM1 and CM2 domains but no PReM domain has yet been identified (Yoo et al., 2025). Nevertheless, CDK5RAP2 contains multiple coiled-coil regions, several of which contain potential Polo/PLK1 phosphorylation sites, which could function in an analogous manner to Cnn-PReM.

**Figure 1.**
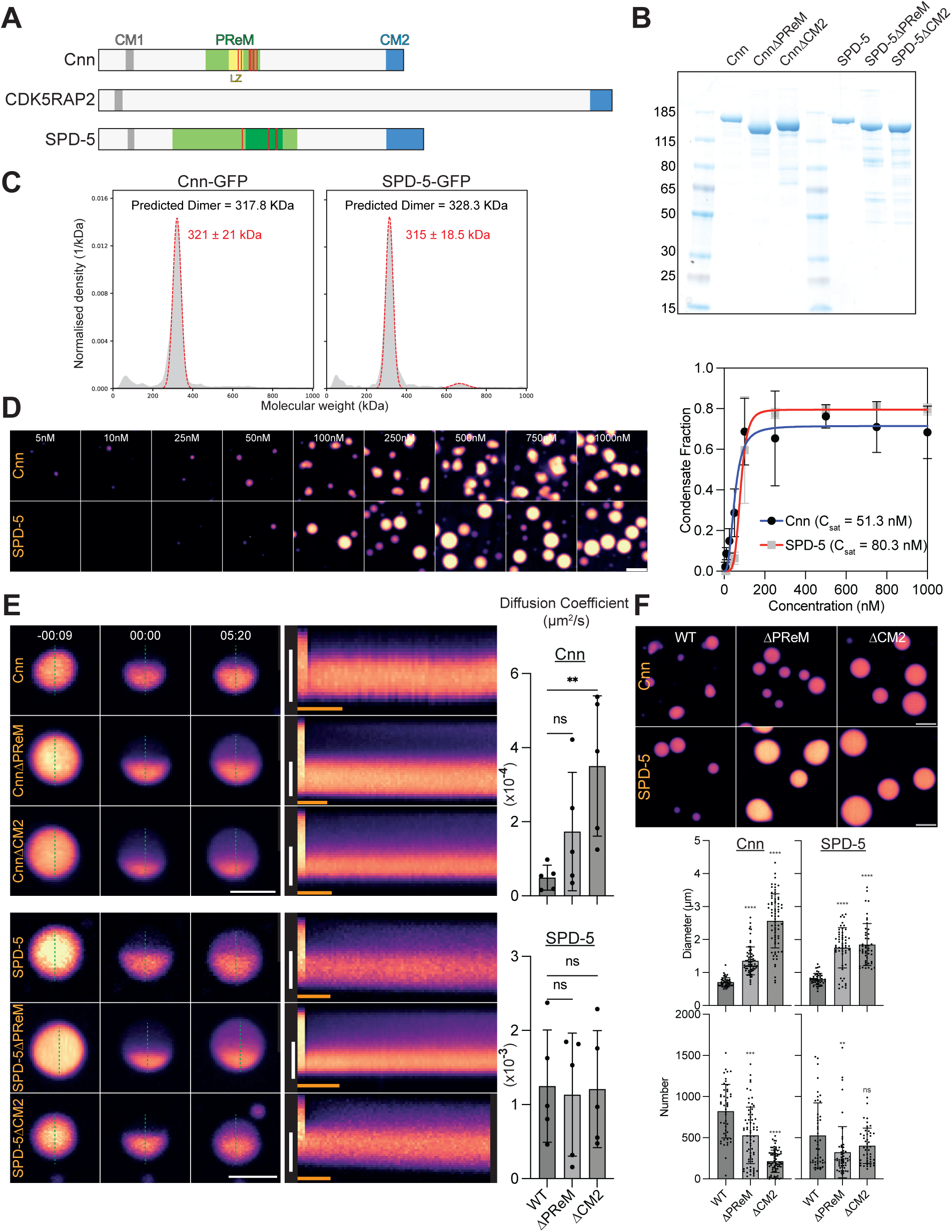
Analysis of full length Cnn and SPD-5 behaviour *in vitro*. **(A)** Schematic illustration of Cnn, CDK5RAP2 and SPD-5 highlighting the position of the CM1 (grey) and CM2 (blue) domains in Cnn and CDK5RAP2 and their proposed equivalents in SPD-5. The Cnn PReM (green, with central LZ in yellow) and putative SPD-5 PReM (light green defined by (Nakajo et al., 2022), dark green defined by (Rios et al., 2024), the latter of which was used in all our studies) are also highlighted. No PReM equivalent has been identified in CDK5RAP2. Red lines indicate functionally important phosphorylation sites in Cnn (Conduit et al., 2014a) and SPD-5 (Woodruff et al., 2015; Ohta et al., 2021). **(B)** Coomassie stained gel of the purified Cnn and SPD-5 proteins analysed here. **(C)** A trace from a Mass photometry experiment showing the MW of the main species of full length Cnn-GFP and SPD-5-GFP. A Gaussian model was fitted to obtain the molecular weight of the dimer (red dotted lines). **(D)** Images show the condensates formed by full length Cnn-GFP and SPD-5-GFP at different concentrations; graph quantifies the fraction of each protein within the condensate at each concentration. A “Specific binding with Hill slope” equation (GraphPad Prism) was fitted, allowing the calculation of *Csat*. **(E)** Images (left) and kymographs (right) from an experiment where the indicated condensates were half photobleached; graphs show calculated diffusion coefficients of fluorescent molecules moving across the bleached zone (mean±SD). **(F)** Images show, and graphs quantify the number and size of, the condensates formed by full length WT, Δ PReM and ΔCM2 versions of Cnn and SPD-5 at 2μM. Scale Bar (D) = 2μm; (E) = 2μm (vertical), 50secs (horizontal).

SPD-5 has no CM1, PReM or CM2 domains that are recognisable by sequence homology, but recent studies have identified domains in SPD-5 that appear to have analogous functions: an N-terminal region required to recruit γ-tubulin complexes (that shares some limited sequence homology with CM1) (Ohta et al., 2021), and both an internal phosphorylated region and a C-terminal region that promote SPD-5 scaffold assembly (potentially analogous to the PReM and CM2 domains, respectively) (Ohta et al., 2021; Nakajo et al., 2022; Rios et al., 2024) (Figure 1A). Thus, the internal interactions that promote Cnn and SPD-5 scaffold assembly may be similar. On the other hand, studies have suggested that the Cnn scaffold is much less dynamic than the SPD-5 scaffold (Feng et al., 2017; Woodruff et al., 2017), perhaps reflecting differences in their physical properties. Here, we investigate how the PReM and CM2 domains of fly Cnn and their potential analogues in worm SPD-5 contribute to scaffold assembly.

## Results

### Full-length Cnn and SPD-5 can form similar condensates but with different dynamic behaviours *in vitro*

To compare the behaviour of full-length Cnn and SPD-5 *in vitro,* we expressed and purified both proteins using a baculovirus/insect-cell FlexiBac expression system (Lemaitre et al., 2019) (Figure 1B). Mass photometry revealed that both purified proteins behaved predominantly as dimers in solution, even at concentrations as low as 10nM (Figure 1C). In the presence of crowding agents full-length SPD-5 has been shown to form spherical condensates (Woodruff et al., 2017), and we found that full-length Cnn exhibited a similar behaviour (Figure 1D). To probe the dynamic behaviour of these condensates, we performed half-bleach experiments and quantified molecular diffusion across the bleached boundary (Figure 1E). SPD-5 condensates were ∼25-fold more dynamic than Cnn condensates (diffusion coefficients of 1.2±0.8×10^-3^ and 4.9±3.4×10^-5^ μm^2^/s, respectively), consistent with previous reports that the Cnn scaffold is less dynamic than the SPD-5 scaffold (Feng et al., 2017; Woodruff et al., 2017).

### CM2 and PReM are not essential for Cnn or SPD-5 condensate assembly *in vitro*

The *Drosophila* Cnn-CM2 and Cnn-PReM domains are essential for efficient Cnn-scaffold assembly *in vivo* (Conduit et al., 2014a; Feng et al., 2017), and potentially analogous domains required for efficient SPD-5 scaffold assembly *in vivo* have been identified in *C. elegans* (Ohta et al., 2021; Nakajo et al., 2022; Rios et al., 2024) (Figure 1A). For simplicity we hereafter refer to these SPD-5 domains as CM2 (SPD-5_1061-1198_) (Nakajo et al., 2022) and PReM (SPD-5_541-677_) (Rios et al., 2024).

To test whether these domains contribute to condensate assembly *in vitro*, we expressed and purified Cnn and SPD-5 proteins in which either CM2 or PReM were individually deleted. For both proteins, the deletion of either domain led to the formation of fewer condensates that were larger in size; an effect that was more pronounced for Cnn than for SPD-5 (Figure 1F). Half-bleach experiments suggested a trend toward higher diffusion coefficients in Cnn condensates lacking either domain, but particularly ΔCM2 (diffusion coefficient of 3.5±1.9×10^-4^ μm^2^/s compared to 4.9±3.4×10^-5^ μm^2^/s for WT) whereas the dynamics of SPD-5 condensates were not significantly altered (Figure 1E)—consistent with previous data that SPD-5 condensation *in vitro* is driven largely by its multiple dispersed coiled-coil domains (Rios et al., 2024). We conclude that the CM2 and PReM domains influence, but are not strictly required for, Cnn or SPD-5 condensate assembly *in vitro*.

### The Cnn PReM domain forms a double helical hairpin

As these domains are important for Cnn and SPD-5 scaffold assembly *in vivo* we wanted to better understand how they contribute to mitotic PCM scaffold assembly. We first focused on *Drosophila* Cnn as, although the full structure of the original PReM domain (Cnn_403-608_) is unknown, this domain contains an internal leucine-zipper (LZ) dimer (Cnn_490-544_) whose crystal structure, in a tetrameric complex with a CM2 dimer, had been solved (Figure 2A) (Feng et al., 2017). AlphaFold3 (AF3) (Abramson et al., 2024) predicted that the Cnn region containing the PReM domain was composed of several helical regions (Figure 2B). The first helix (H1; Cnn_377-443_) was not predicted to interact with the other helices, and it extended N-terminally to Alanine 377, meaning that the original PReM (Cnn_403-608_) (Feng et al., 2017) contained only approximately half of H1. The second helix (H2; Cnn_T462-K544_) contained the original LZ that interacts with CM2 (Cnn_490-544_—highlighted in *yellow*), which was followed by two shorter helices: H3 (Cnn_A554-T583_) and H4 (Cnn_S589-601_ or Cnn_S587-607_ depending on the precise input sequence used for the prediction). These 3 helices (H2-H4) were predicted to form a structural domain, with H3 and H4 folding back to interact with each other and with H2 to form a double helical hairpin (Figure 2B). AF3 predicted a very similar helical organisation for a potential PReM dimer (Figure S1). Although this is not a very confident dimer prediction (ipTM = 0.4), the PReM domain is almost certainly dimeric, as all PReM-related constructs we have made to date are dimeric in solution (Figure S2), as is full length Cnn (Figure 1C). For simplicity, however, we show the PReM as a monomer in some schematics to better illustrate the boundaries of the deletion constructs we tested here.

**Figure 2.**
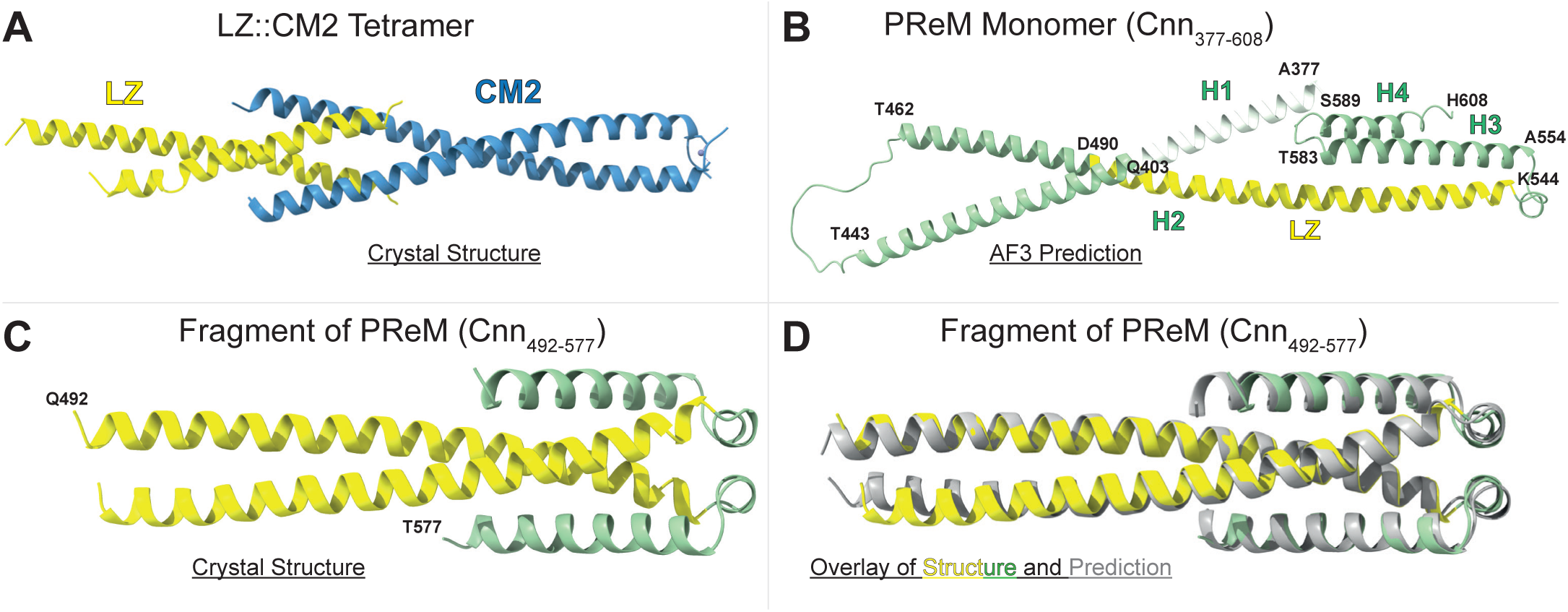
The *Drosophila* PReM domain adopts a helical hairpin fold. Ribbon representations of various PReM domain crystal structures or AF3 predictions. **(A)** Previously published crystal structure of the PReM LZ::CM2 interaction. **(B)** AF3 prediction of the Extended PReM monomer. The PReM is predicated to comprise 4 helices (H1-H4) with H3 and H4 helices folding back on each other and on the C-terminal region of H2 to form a helical hairpin. The original PReM domain is coloured green (except for the internal LZ domain, yellow). AF3 predicts that the N-terminal boundary of the original PReM (Q403) lies within a helix that extends to A377 (white). **(C,D)** Crystal structure (C) and overlay of the crystal structure (yellow/green) with an AF3 prediction (grey) (D) of a fragment of the PReM domain containing the LZ (C-terminal region of H2) and most of H3. This fragment forms a dimer *in crystallo*, confirming the predicted helical-hairpin fold of this part of the PReM domain.

We attempted to validate this helical hairpin structure by crystallising various PReM constructs but did not obtain any useful data with any constructs that contained both helices H1 *and* H2 or both helices H3 *and* H4, potentially because H1 and H4 exhibit flexibility in their packing against a more stable H2/H3 core. In support of this possibility, we obtained crystals of a shorter PReM dimer fragment (Cnn_490-579_) that lacked H1 and H4, but that contained most of H2 together with H3 (Figure 2C). These crystals diffracted to 1.96Å and the crystal structure aligned well with the AF3 dimer prediction (ipTM=0.73; 86 Cα pairs with rmsd = 0.910Å) confirming the overall dimeric helical hairpin structure of the core H2/H3 helices (Figure 2D).

### Defining the PReM domain regions required for scaffold assembly *in vitro* and *in vivo*

We previously showed that the LZ::CM2 interaction is crucial for the assembly of the PReM::CM2 scaffold *in vitro* and the endogenous Cnn scaffold *in vivo* (Feng et al., 2017). Moreover, the phosphorylation of the more C-terminal region of the PReM (that forms H3 and H4 of the dimeric helical hairpin) is also important for scaffold assembly *in vitro* and *in vivo* (Conduit et al., 2014a; Feng et al., 2017). It was not clear, however, whether the PReM regions N-terminal to the LZ are required for these processes. To test this, we first purified an N-terminally extended version of the PReM that contains all of helix 1 (H1) (Cnn_377-608_, or Extended-PReM). We found that monomeric-NeonGreen (NG) fusions to both the Original-PReM (Cnn_403-608_) and Extended-PReM formed macromolecular scaffolds when mixed with CM2 *in vitro* (Figure 3A,B,E). This was also the case if H1 was entirely deleted from the PReM construct (Cnn_462-608_, or PReMΔH1) (Figure 3C,E). If, however, we additionally deleted the N-terminal ∼1/3 of H2 (Cnn_490-608_, or PReMΔH1ΔNTH2) (leaving only the LZ region of H2 intact), macromolecular complexes with CM2 could still form, but these appeared to be large particles that tended to bunch together on the microscope slide, rather than extended scaffolds (Figure 3D,E). These Cnn_490-608_::CM2 particles had an average mass of ∼1MDa (Figure S3A) and they could be visualised by negative stain EM (Figure S3B), but they lacked any obvious repetitive structure and were not suitable for Cryo-EM analysis. We conclude that H1 is not essential for PReM::CM2 scaffold assembly *in vitro*, but that deletions that remove the N-terminal part of H2 start to perturb scaffold assembly even if the LZ and adjacent helical hairpin are still intact (although some large, but more discreet, macroscopic assemblies can still form).

**Figure 3.**
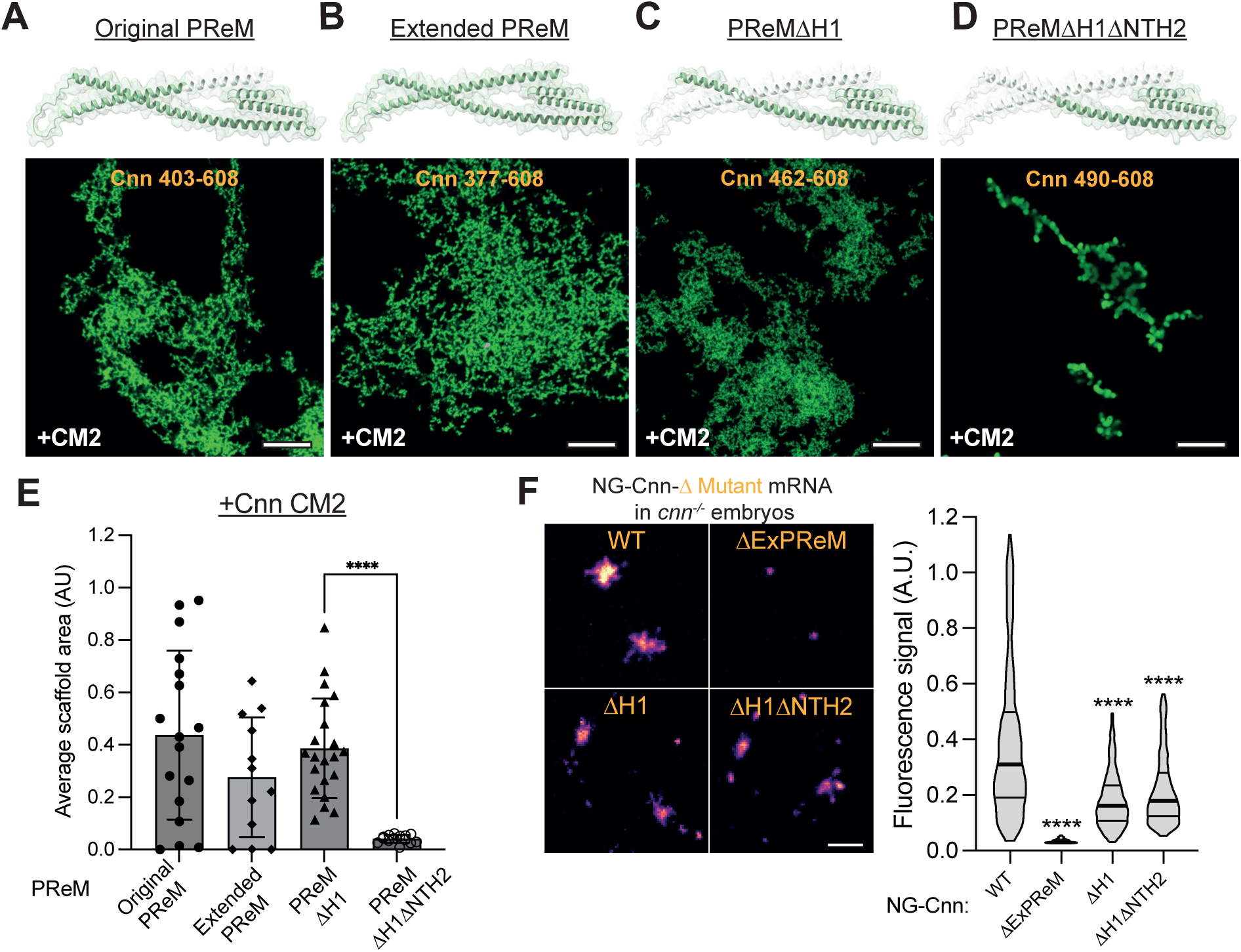
Analysis of PReM regions required for scaffold assembly *in vitro* and *in vivo*. **(A-D)** Ribbon representations illustrate the PReM domain constructs being tested here (tested region highlighted in green). Images below each construct show a typical microscope field of the structures observed when NG-fusions of each construct were mixed with CM2 *in vitro*. **(E)** Graph quantifies the average area of NG-structure observed by microscopy for each construct shown in (A-D). **(F)** Images illustrate, and violin plot quantifies, the in vivo centrosomal fluorescence intensity (bars indicate mean and first and third quartile) of each protein when mRNA encoding a NG-fusion of each construct was injected into embryos laid by *cnn^-/-^* females (hereafter *cnn^-/-^* embryos) that lack endogenous Cnn. Statistical significance was assessed using an unpaired t test in GraphPad Prism (****p < 0.0001). Scale bar (A-D) = 10μm; (F) = 2μm.

To assess whether H1 was required for Cnn scaffold assembly *in vivo*, we injected mRNA encoding either WT or H1-deleted Cnn (CnnΔH1) into embryos laid by *cnn^-/-^* mothers that lacked any endogenous Cnn protein (hereafter *cnn^-/-^* embryos). As expected, the deletion of the entire extended PReM (NG-CnnΔExPRem) strongly suppressed scaffold assembly *in vivo*, but the deletion of only H1 (CnnΔH1) had a much milder, although still significant, effect (Figure 3F). Interestingly, when we deleted the first ∼1/3 of H2 in addition to deleting H1 (CnnΔH1ΔNTH2), the ability to form a Cnn scaffold was still only mildly perturbed—to about the same extent as deleting only H1 (Figure 3F). Together with published data (Conduit et al., 2014a; Feng et al., 2017), these data indicate that the C-terminal helical hairpin region of the PReM is particularly important for physiological Cnn scaffold assembly.

### The PReM helical hairpin appears to autoinhibit Cnn scaffold assembly

The helical hairpin that lies C-terminal to the original LZ (Cnn_490-544_) region contains 8 Serines that are highly conserved in *Drosophila* species and most of which match the minimal PLK1 consensus phosphorylation sequence reasonably well (N/D/E-X-S-Φ, where S is the phosphorylated Serine and Φ indicates a hydrophobic amino acid) (Nakajima et al., 2003; Santamaria et al., 2011) (Figure 4A). *In vivo*, mutating combinations of these Serines to Alanines perturbs Cnn scaffold assembly to varying degrees, suggesting that multiple Serines within this region are phosphorylated to promote Cnn scaffold assembly (Conduit et al., 2014a). We therefore focused on trying to understand how phosphorylation might influence the helical hairpin structure.

**Figure 4.**
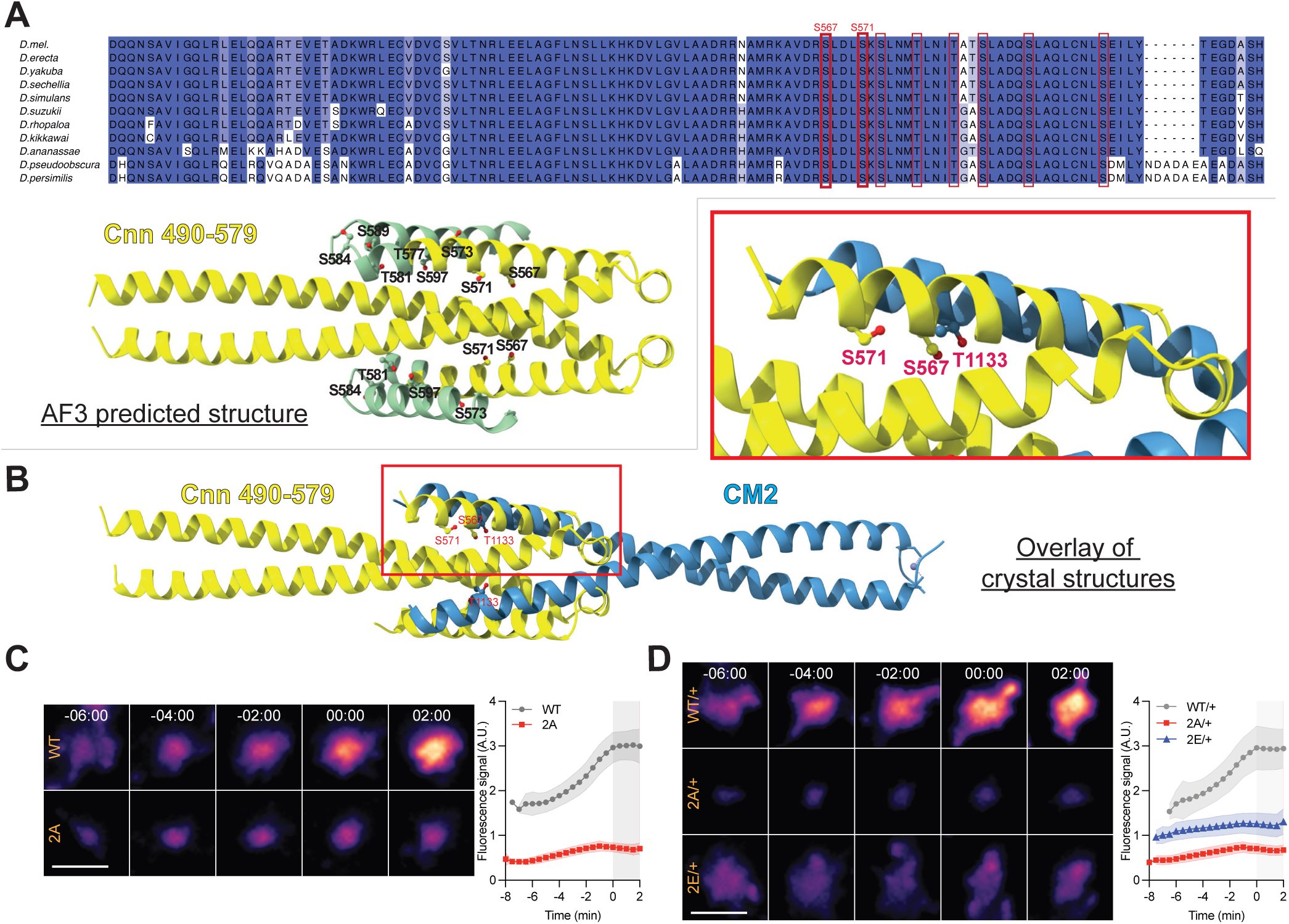
The PReM hairpin blocks CM2 binding, but this appears to be relieved by the phosphorylation of S567 and/or S571. **(A)** Multiple Sequence Alignment of the PReM domain from several *Drosophila* species showing the positions of several conserved Ser/Thr’s (red boxes) in the helical hairpin region (highlighted in the ribbon diagram of the AF3 predicted structure below). **(B)** An overlay of the crystal structures of the PReM helical hairpin (containing the LZ and most of H3), and the CM2 domain when bound to the LZ (only CM2 shown here, aligned to the LZ in the PReM helical hairpin structure). The binding of H3 to H2 within the PReM occludes the binding site on H2 for the CM2 domain. Inset shows the position of S567 and S571 within the H2::H3 PReM interface and of T1133 in the PReM::CM2 interface. **(C,D)** Images illustrate, and graphs quantify, the centrosomal fluorescence intensity of WT Cnn-NG or Cnn-2A-NG expressed in *cnn^-/-^* embryos (C) or WT Cnn-NG or Cnn-2A-NG or Cnn-2E-NG laid by *cnn^+/-^* heterozygous females (so the embryos contain some endogenous unlabelled WT Cnn) (D) during nuclear cycle 12. Graphs illustrate the mean±SD

Overlaying our crystal structure of the PReM helical hairpin (Cnn_490-579_), with our previous crystal structure of the PReM LZ bound to CM2 revealed that the folding-back of H3 occluded the normal binding site for CM2 on H2 (Figure 4B). Interestingly, S567 in H3 occupied a very similar position in the crystal structure to the highly conserved T1133 of CM2 when bound to H2 (*inset*, Figure 4B). This was potentially informative, as a single T1133E substitution essentially abolishes the PReM::CM2 interaction *in vitro* as well as Cnn scaffold assembly *in vivo*, while phospho-specific antibodies reveal that Cnn-S567 seems to be phosphorylated only on Cnn molecules that have recently become incorporated into the Cnn scaffold (Feng et al., 2017). This immediately suggested the hypothesis that the PReM H2/H3 hairpin normally prevents CM2 binding, and that phosphorylation of S567 (and perhaps also the nearby S571, which is also buried in the helical hairpin interface—*inset*, Figure 4B) “opens up” the hairpin to allow CM2 to bind H2 and so allow Cnn molecules phosphorylated in this way to participate in scaffold assembly.

To test this possibility, we mutated S567 and S571 to Alanine (Cnn-2A) and generated transgenic lines that expressed NG-Cnn-2A in *cnn^-/-^* embryos. As reported previously (Wong et al., 2022, 2024), WT-NG-Cnn levels started to accumulate at centrosomes in the run-up to mitosis, and levels peaked during mitosis (mitosis is indicated by the grey area in the graph in Figure 4C, starting at t=0). In contrast, although NG-Cnn2A still localised to centrosomes, it could not assemble a robust scaffold (Figure 4C), consistent with our hypothesis that S567 and/or S571 phosphorylation helps to open the PReM helical-hairpin to allow CM2 binding and scaffold assembly.

Importantly, PReM can bind CM2 *in vitro* even when it has not been phosphorylated, suggesting that, at least in this *in vitro* context, the helical-hairpin can “breath” to allow CM2 binding even in the absence of phosphorylation (Feng et al., 2017). To our surprise, however, we could detect no interaction between CM2 and PReM-2A_462-608_ or PReM-2A_490-608_ *in vitro* (Figure 5A), and these PReM constructs could no longer form scaffold structures when mixed with CM2 (Figure 5B). This result questions our interpretation that the inability of GFP-Cnn-2A to form a scaffold *in vivo* is due to a failure to phosphorylate S567 and/or S571 to open-up the PReM for CM2-binding; these S/A substitutions seem to render the PReM domain unable to bind CM2 at all.

**Figure 5.**
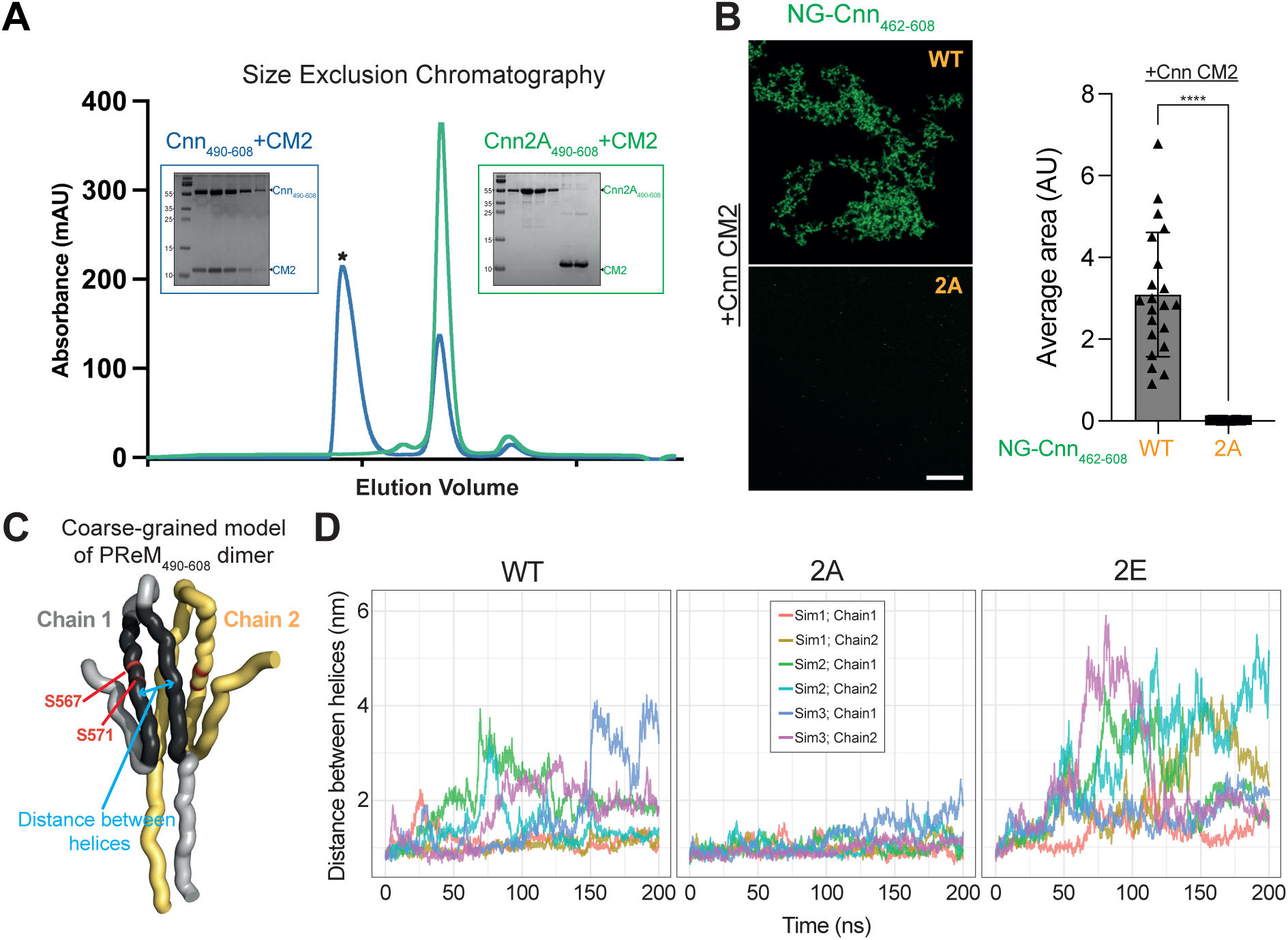
The Cnn-2A mutations block the PReM::CM2 interaction and appears to perturb the “breathing” of the PReM helical hairpin. **(A)** Panel shows the trace from a size-exclusion chromatography experiment (insets show the accompanying SDS-gels) illustrating the behaviour of the WT (blue) and 2A-mutant (green) PReM domains when mixed with CM2 in vitro. WT PReM forms high-MW complexes with CM2 that elute in the void volume (asterisk) whereas the PReM Cnn-2A mutant does not bind to CM2. **(B)** Images show a typical microscope field of the structures observed when NG-fusions of WT or 2A PReM were mixed with CM2 *in vitro*; the bar-chart quantifies the average area of scaffold observed by microscopy for each PReM construct (mean±SD). Statistical significance was assessed using an unpaired t test in GraphPad Prism (****p < 0.0001). Scale bar = 10μm **(C)** Schematic illustration of the coarse-grain model of the PReM (Cnn_490-608_) helical hairpin dimer, highlighting the position of S567 and S571 and the point where the distance between the H2 and H3 helices in the hairpin structure were measured during molecular simulations. **(D)** Graphs plot the distance between the H2 and H3 helices during 3 separate simulations of 200ns. This generated 6 traces as the distance between the helices was calculated for both chains of the dimer independently.

To try to understand the molecular basis for this surprising result, we first tested whether the 2A mutation disrupted the helical hairpin fold, but we found no appreciable difference in the crystal-structure of Cnn-2A_490-579_ when compared to WT Cnn_490-579_ (81 Cα pairs with rmsd = 0.718 Å; Figure S4). We wondered, therefore, whether the 2A substitutions might lock the hairpin into a more stable conformation that can no longer breath to allow CM2 binding. To assess this possibility, we performed coarse-grained molecular simulations with WT and 2A versions of PReM_490-608_, quantifying the breathing of the two main helices (H2 and H3) by calculating the distance between their centres at each timepoint of the simulation (Figure 5C). This revealed that the WT protein exhibited significant breathing in the simulation, which was dramatically suppressed by the 2A substitutions (Figure 5D). We also ran simulations with an S567E, S571E (2E) mutant protein, mimicking the phosphorylation of S567 and S571. This increased the breathing of the hairpin, supporting our hypothesis that phosphorylation “opens” the PReM hairpin to promote CM2 binding (Figure 5D).

We also attempted to examine the localisation of a full length GFP-Cnn-2E in *cnn^-/-^* embryos but found that the expression of this protein was dominant male sterile. This meant that we could not generate flies expressing this protein in a homozygous *cnn^-/-^* background. We therefore compared the behaviour of WT NG-Cnn, NG-Cnn-2A and NG-Cnn-2E in the presence of a single copy of the endogenous (untagged) Cnn gene—and thus presumably in the presence of some WT:2A or WT:2E Cnn heterodimers. In this system, NG-Cnn-2E exhibited an intermediate phenotype, incorporating into a Cnn-scaffold slightly better than the embryos expressing NG-Cnn-2A, but not as well as the embryos expressing WT NG-Cnn (Figure 4D). This suggests that although the Cnn-2E substitutions should theoretically open the PReM domain to allow CM2 binding, Cnn-2E cannot form a normal scaffold.

### The CM2 domain of human CDK5RAP2, but not worm SPD-5, can partially substitute for the fly CM2 domain

These studies emphasise the importance of the PReM::CM2 interaction for Cnn-scaffold assembly in flies. As discussed above, the sequence of the C-terminal CM2 domain is relatively well conserved between flies and mammals (∼22% identity over 69aa; Figure S5), and AF3 predicts that the fly-Cnn and human-CDK5RAP2 CM2 domains adopt a similar fold (Figure 6A). The SPD-5 CM2 domain, however, exhibits very little sequence conservation with the fly and human proteins (∼4% identity over 69aa; Figure S5) and AF3 predicted that this region adopted a different helical architecture (Figure 6A). Strikingly, when we purified NG-fusions of the fly, human and worm CM2 proteins and mixed them individually with the *Drosophila* PReM_462-608_ domain, the fly and human CM2 domains formed similar PReM::CM2 macromolecular networks, but worm CM2 did not (Figure 6B).

**Figure 6.**
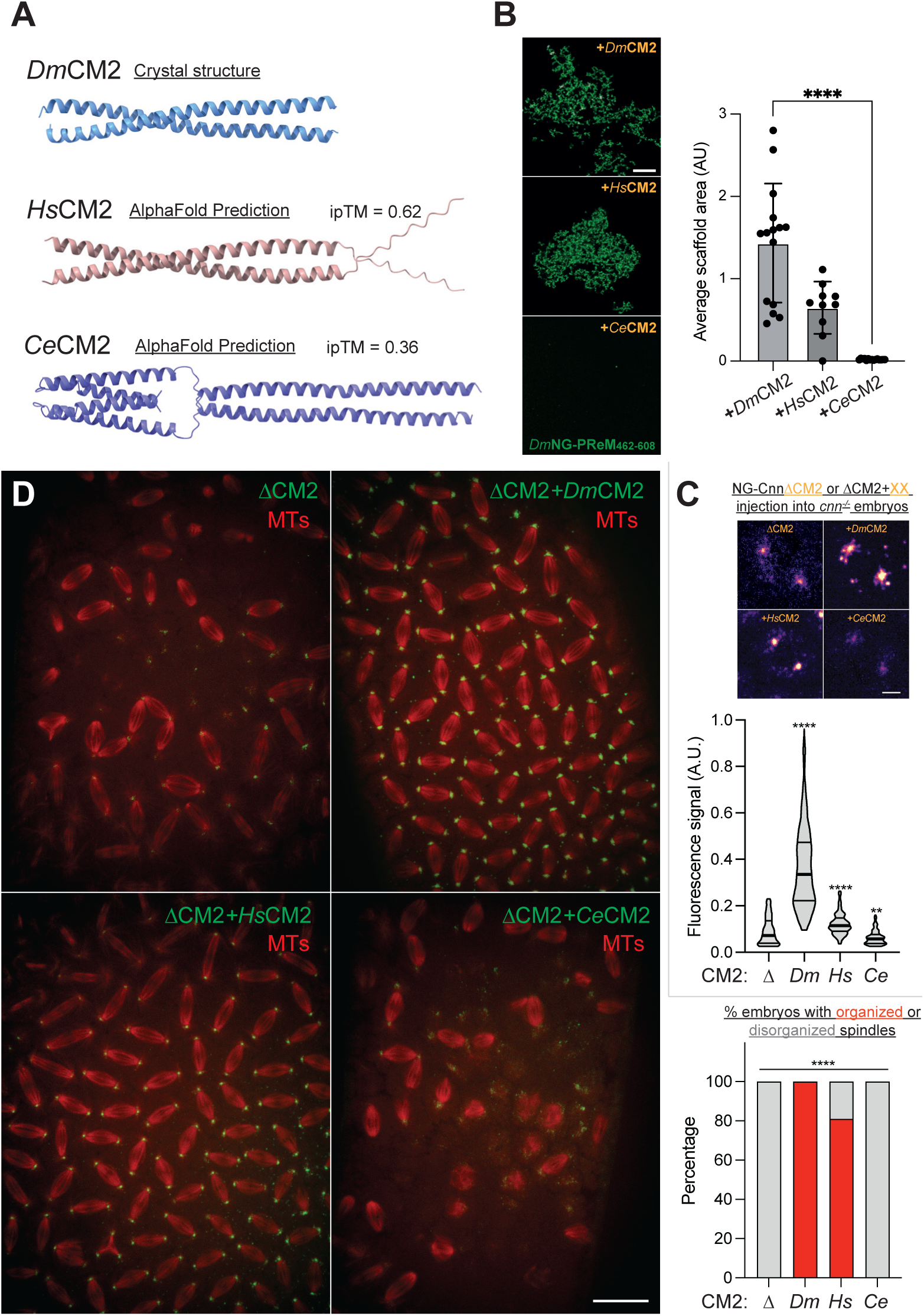
Human CDK5RAP2 CM2 can partially substitute for *Drosophila* Cnn CM2 in scaffold assembly. **(A)** Crystal structures or AF3 predictions of *Drosophila*, Human or *C. elegans* CM2 dimers. **(B)** Images show a typical microscope field of the structures observed when fly (*Dm*), Human (*Hs*) or worm (*Ce*) CM2 were mixed with fly NG-PReM *in vitro*; the bar-chart quantifies the average area of scaffold observed by microscopy for each CM2 construct (mean±SD). Statistical significance was assessed using an unpaired t test in GraphPad Prism (****p < 0.0001). **(C)** Images illustrate, and violin plot quantifies, the *in vivo* centrosomal fluorescence intensity (bars indicate mean and first and third quartile) of each protein when mRNA encoding a NG-fusion of each construct was injected into *cnn^-/-^* embryos. Statistical significance was assessed using an unpaired t test in GraphPad Prism (****p < 0.0001). **(D)** Images illustrate, and bar chart quantifies, the ability of chimeric full length Cnn molecules to rescue the embryonic defects in *cnn^-/-^* embryos. These embryos exhibit a range of mitotic defects that were completely rescued by the injection of mRNA encoding Cnn containing the *Drosophila* CM2 domain (100% rescue, N=16), partially rescued by Cnn containing the Human CM2 domain (80%, N=16), but were not detectably rescued by Cnn containing the worm CM2 domain (0%, N=16). Images of these embryos were randomised by one worker and analysed blind by another worker who had not previously seen any of the images but was familiar with the *cnn^-/-^* embryo phenotype. Contingency significance (i.e., the significance of the difference in the proportion between the two groups) was calculated using a Fisher’s exact test (****P < 0.0001). Scale bar (A-D) = 10μm; (F) = 20μm.

To test whether the human CM2 domain could substitute for the fly CM2 domain *in vivo*, we injected chimeric versions of NG-Cnn in which we substituted the human or worm CM2 domains for the fly CM2 domain. The fly/human chimera localised to centrosomes (although not as strongly as the WT protein) (Figure 6C) and it rescued the centrosome and associated developmental defects in *cnn^-/-^* embryos very well (Figure 6D), whereas the fly/worm chimera did not (Figure 6C,D). Taken together, these experiments suggest that the CM2 domain of fly Cnn and human CDK5RAP2 can cooperate with *Drosophila* PReM to promote the assembly of structurally-related scaffold structures, whereas the CM2 domain of worm SPD-5 cannot.

In support of this conclusion, although the PReM domain of fly Cnn and the putative PReM domain of worm SPD-5 (Rios et al., 2024) appear to adopt helical hairpin structures, these structures are quite different (Figure 7A). Moreover, although both domains contain several Ser/Thr residues whose phosphorylation is clearly important for scaffold assembly, none of the 3 potential phosphorylation sites in the worm protein are located in positions where their phosphorylation would be predicted to “open” the helical hairpin structure (although we note the caveat that the worm SPD-5 PReM structure is a prediction, whereas the positioning of S567 and S571 in the fly PReM has been confirmed by crystallography). We also found no evidence for an *in vitro* interaction between the worm SPD-5 PReM and CM2 domains (Figure 7B) and, when mixed, NG-fusions of these domains did not detectably form scaffold structures, even after these domains had been phosphorylated *in vitro* by PLK-1 (Figure 7B,C).

**Figure 7.**
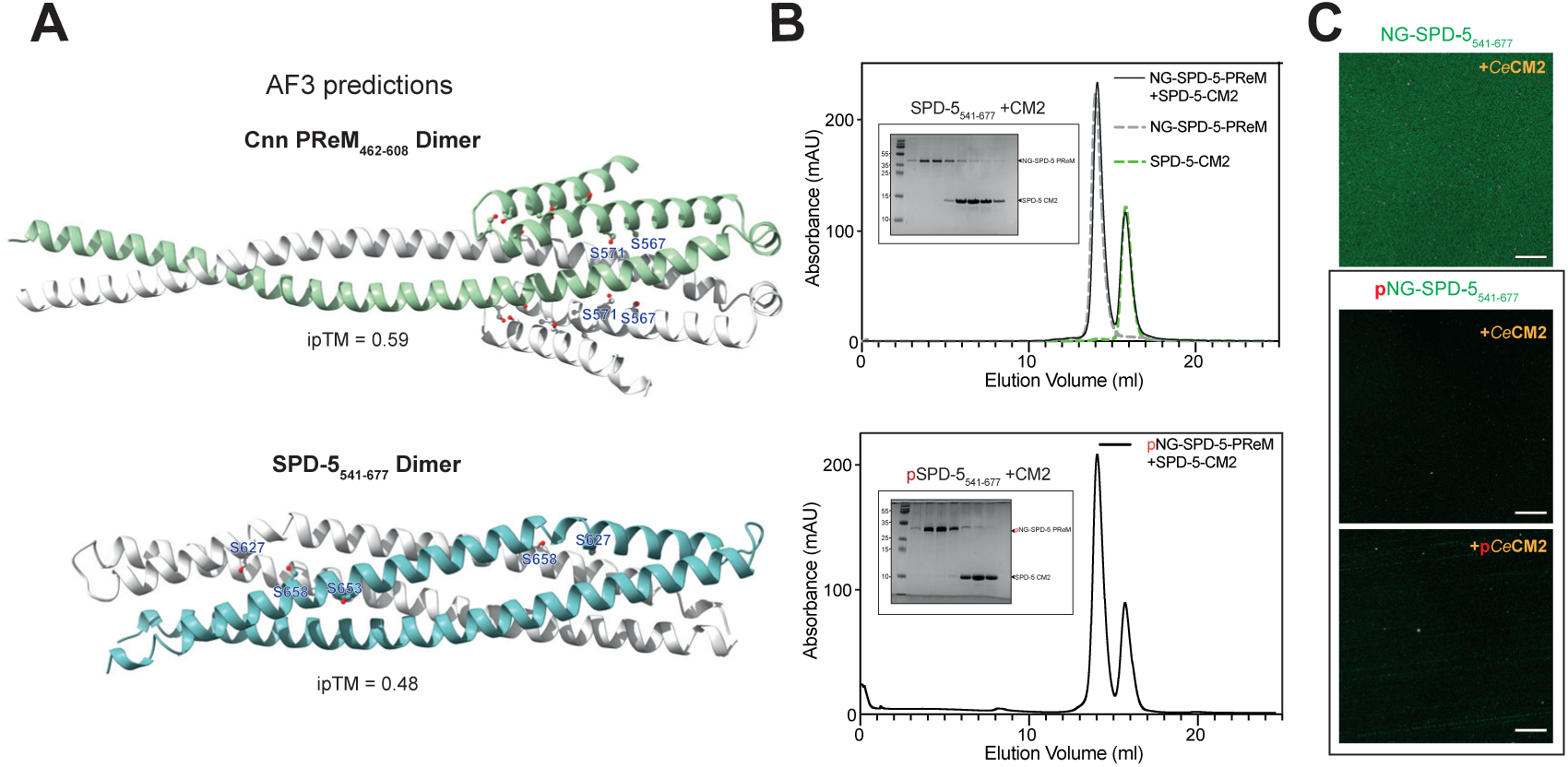
The putative PReM and CM2 domains in C.elegans SPD-5 do not detectably interact. **(A)** AF3 predictions of *Drosophila* or *C. elegans* PReM dimers. The putative SPD-5 PReM helical hairpin dimer was identified and shown to be important for SPD-5 scaffold assembly previously (Rios et al., 2024). **(B)** Panels show traces from size-exclusion chromatography experiments (insets show relevant SDS-gels) illustrating the behaviour of SPD-5-CM2, NG-SPD-5-PReM, or a mixture of the two (black trace and gel inset), either without phosphorylation or when NG-SPD-5-PReM has been phosphorylated by PLK1 kinase prior to mixing (as phosphorylation by PLK1 stimulates the ability of the SPD-5-putative PReM to form scaffolds *in vitro* and *in vivo*) (Rios et al., 2024). **(C)** Images how typical microscope fields observed when NG-SPD-5-PReM (non-phosphorylated, or phosphorylated by PLK1) were mixed with CM2 (non-phosphorylated or phosphorylated by PLK1) *in vitro*. (C) Graph quantifies the average area of scaffold observed by microscopy for each construct shown. Scale bar = 10μm

Finally, as the human CDK5RAP2 CM2-domain can partially function in place of the fly Cnn CM2 domain, we wondered if we could identify the CDK5RAP2 equivalent of the PReM domain. We used AF3 to predict the plausibility of interactions between human CDK5RAP2-CM2 and all the other predicted coiled-coil domains within CDK5RAP2. No high-confidence interactions were identified (ipTM score >0.5) but the highest confidence prediction was structurally somewhat reminiscent of the Cnn-PReM::CM2 interaction and had an ipTM score of 0.46 (Figure 8A). Moreover, in isolation, this putative PReM domain (CDK5RAP2_535-639_) was predicted to adopt a structure somewhat reminiscent of the Cnn-PReM helical hairpin, albeit with low-confidence (ipTM = 0.17). It also contained several Ser/Thr residues, although none of these were buried in the helical hairpin interface (again with the caveat that this is a relatively low confidence prediction, not an experimentally determined structure) (Figure 8B). Intriguingly, when mixed with purified human CM2, a NG-fusion of this putative PReM domain formed large macromolecular structures that were reminiscent of those formed by the equivalent fly proteins (Figure 8C). Previous attempts to identify the CDK5RAP2 PReM domain did not identify any regions of CDK5RAP2 that were able to do this (Yoo et al., 2025). Thus, CDK5RAP2_535-639_ is a plausible candidate for the CDK5RAP2 PReM domain.

**Figure 8.**
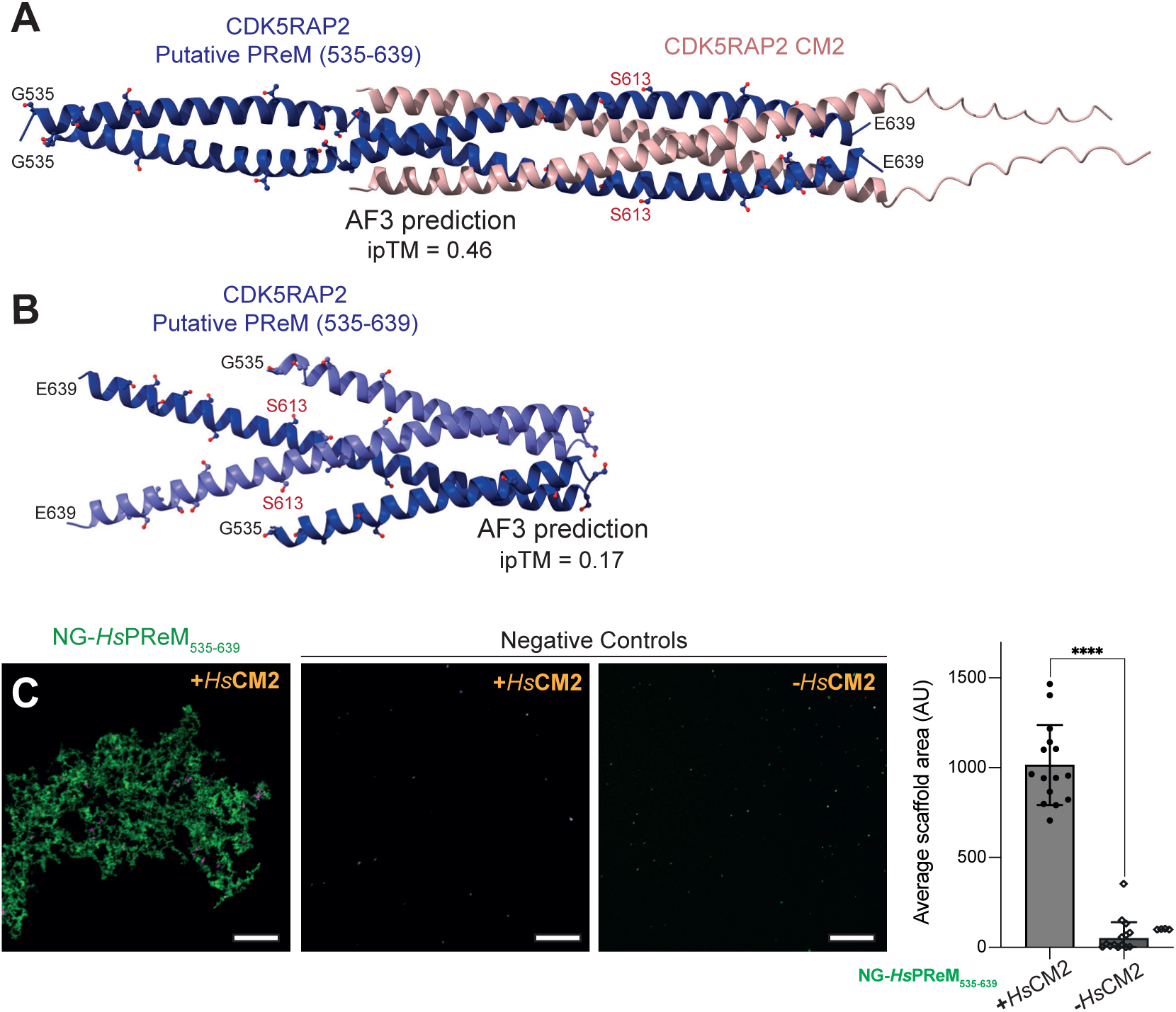
Identification of a PReM domain candidate in Human CDK5RAP2. **(A,B)** AF3 predictions of a potential PReM domain helical hairpin dimer in Human CDK5RAP2 (A), and its potential interaction with the Human CM2 domain (B). **(C)** Images show a typical microscope field observed when NG-CDK5RAP2-PReM was mixed with CDK5RAP2-CM2 in vitro. Graph quantifies the average area of scaffold observed by microscopy for each condition shown. Scale bar = 10 μm

## Discussion

The dramatic expansion of the PCM in preparation for mitosis seems to be a universal feature of centrosome behaviour in the diverse range of species and cell types that use centrosomes to help guide cell division. Flies and worms use a similar set of proteins (Spd-2/SPD-2, Polo/PLK-1 and Cnn/SPD-5) to assemble a scaffold structure that gives mechanical strength to the expanded PCM and ultimately recruits most of the other mitotic PCM components. Although proteins within the Spd-2/SPD-2 family and within the Polo/PLK1 family are clear homologues, Cnn and SPD-5 share no obvious sequence homology. Here we have tried to establish whether the internal interactions that promote Cnn and SPD-5 scaffold assembly are similar.

### Cnn and SPD-5 form condensates *in vitro* but with distinct dynamic properties

We first compared the behaviour of purified recombinant full-length Cnn and SPD-5 *in vitro*. As shown previously for SPD-5 (Woodruff et al., 2017), both proteins can form spherical condensates in the presence of crowding agents. Half-FRAP experiments revealed a striking difference in their internal dynamics: SPD-5 condensates were ∼25-fold more dynamic than those formed by Cnn, indicating that SPD-5 molecules exchange more rapidly within condensates than Cnn molecules. Interestingly, however, the deletion of either the PReM or CM2 domains did not prevent condensate formation *in vitro* for either protein, although these deletions had distinct effects on dynamics. For Cnn, the deletion mutants formed fewer but larger condensates that were more dynamic than full-length Cnn, whereas the dynamics of the SPD-5 deletion condensates were not significantly affected. Our observation that Cnn and SPD-5 condensates still form *in vitro* in the absence of these domains implies that other, weaker, interactions can support self-assembly under the buffer conditions used in the assay. These interactions may be physiologically relevant, but not strong enough to support PCM assembly *in vivo*. Alternatively, they may be non-physiological and arise from the buffer conditions and crowding agents used in the assay. Further work will be needed to determine which features of the in vitro condensates reflect the organisation of the PCM scaffold in vivo.

### The role of the PReM and CM2 domains in promoting scaffold assembly

An important insight from our work is that the *Drosophila* PReM appears to form an auto-inhibited helical hairpin that in its “closed” configuration cannot bind CM2. Our data suggests that the phosphorylation of one or both of two conserved Serines (S567 and S571) buried in the helical hairpin interface can “open” the helical hairpin to allow CM2 binding and scaffold assembly. This potentially explains our previous observation that S567 appears to be phosphorylated on Cnn molecules that have newly incorporated into the Cnn scaffold (Conduit et al., 2014a). Mutating S567 and S571 to phospho-mimicking Glu only partially restores Cnn scaffold assembly *in vivo*, suggesting either that these substitutions do not fully mimic the effects of phosphorylation, or that these Serines may have additional functions in scaffold assembly that are not related to their phosphorylation. The phosphorylation of several other Ser/Thr’s within the helical hairpin contributes to Cnn scaffold assembly (Conduit et al., 2014a), and we show here that the N-terminal region of Helix 2 (that is not directly involved in the formation of the helical hairpin) also promotes scaffold assembly. Thus, the phosphorylation of S567 and/or S571 may be important for initiating Cnn scaffold assembly, but other regions of the PReM domain are also clearly involved in this process. How these other regions promote PReM::CM2 scaffold assembly *in vitro*, and Cnn scaffold assembly *in vivo*, remains to be determined.

### The function of putative PReM and CM2 domains in other scaffolding proteins

To test whether similar principles apply in vertebrates, we examined human CDK5RAP2. Human CDK5RAP2 has a C-terminal CM2 domain and we show here that it can form macromolecular scaffolds when mixed with the *Dm*PReM domain, and that a chimeric Cnn molecule in which *Dm*CM2 had been replaced with *Hs*CM2 can partially restore Cnn function *in vivo*. Moreover, we identified a putative *Hs*PReM domain that can form macromolecular scaffolds when mixed with *Hs*CM2 *in vitro*. Together, these data suggest that human CDK5RAP2 and *Drosophila* Cnn use similar molecular self-interactions to form centrosomal scaffolds. From the predicted structure, however, it is not obvious that *Hs*PReM forms an autoinhibited helical hairpin whose phosphorylation would open the hairpin to allow binding to *Hs*CM2. Thus, the mechanisms regulating the putative *Hs*PReM::*Hs*CM2 interaction may be different in flies and humans. However, the AF3 prediction for *Hs*PReM helical hairpin is low confidence (ipTM = 0.17), so the structural basis of any regulated CDK5RAP2 scaffold assembly in humans remains unclear.

Although putative PReM and CM2 domains have been identified in worm SPD-5, we found no evidence that these form interactions in a manner similar to the equivalent fly domains. We could not find conditions where *Ce*PReM interacts with *Ce*CM2 *in vitro*, and unlike *Hs*CM2, the *Ce*CM2 is unable to form scaffolds with *Dm*PReM *in vitro*, or to partially substitute for *Dm*CM2 domain in full length Cnn scaffold assembly *in vivo*. Thus, although one cannot make definitive conclusions from such negative data, it seems likely that Cnn and SPD-5 use distinct molecular interactions to drive scaffold assembly. Intriguingly, worm SPD-5 does have a recognisable CM1 domain that binds γ-tubulin complexes (Ohta et al., 2021), suggesting at least some evolutionary conservation between SPD-5 and the Cnn/CDK5RAP2 family of proteins.

Together, our findings highlight both conservation and divergence in the molecular mechanism underlying centrosomal scaffold assembly. While flies and humans appear to rely on related PReM-CM2 interactions, that in flies are clearly regulated by phosphorylation, worms appear to have diverged to use distinct mechanisms, although these are clearly also regulated by phosphorylation. These similarities in function, yet apparent differences in molecular interactions, emphasize the evolutionary plasticity of PCM scaffold assembly and the robustness of outcome: a mechanically strong structure capable of recruiting a broad set of proteins to the mitotic PCM.

## Materials and Methods

### Drosophila melanogaster stocks and husbandry

The *Drosophila* stocks used, generated and/or tested in this study are listed in Table 1. The precise stocks used in each experiment (and the relevant Figure) are listed in Table 2. Flies were maintained on *Drosophila* culture medium (0.68% agar, 2.5% yeast extract, 6.25% cornmeal, 3.75% molasses, 0.42% propionic acid, 0.14% tegosept, and 0.7% ethanol) in 8cm x 2.5cm plastic vials or 0.25-pint plastic bottles. For microscopy and immunoblot experiments, flies were placed in embryo collection cages on fruit juice plates (see below) with a drop of yeast paste. Fly handling were performed as previously described (Roberts, 1998).

**Table 1:** *Drosophila* stocks used in this study.

| Allele | Source |
| --- | --- |
| cnn<sup>f04547</sup> | Exelixis stock no. f04547, Exelixis Stock Centre (Harvard Medical School, Boston, MA). |
| cnn<sup>HK21</sup> | (Megraw *et al*, 1999; Vaizel-Ohayon & Schejter, 1999) |
| Ubq-NG-Cnn | (Wong *et al*, 2022) |
| Ubq-NG-Cnn-2A | Generated in this study |
| Ubq-NG-Cnn-2E | Generated in this study |

**Table 2:** *Drosophila* stocks used in specific experiments.

| Genotype | Purpose | Figure |
| --- | --- | --- |
| *cnn*<sup>f04547</sup> / *cnn*<sup>HK21</sup> | For injection of various truncated Cnn mutants | 3F |
| *cnn*<sup>f04547</sup>, Ubq-NG-Cnn / *cnn*<sup>HK21</sup> | Comparing the dynamics of Cnn phosphorylation null mutants | 4C |
| *cnn*<sup>f04547</sup>, Ubq-NG-Cnn-2A / *cnn*<sup>HK21</sup> |  |  |
| Ubq-NG-Cnn / + | Comparing the dynamics of Cnn phosphorylation null and mimicking mutants | 4D |
| Ubq-NG-Cnn-2A / + |  |  |
| Ubq-NG-Cnn-2E / + |  |  |

**Table 3:**
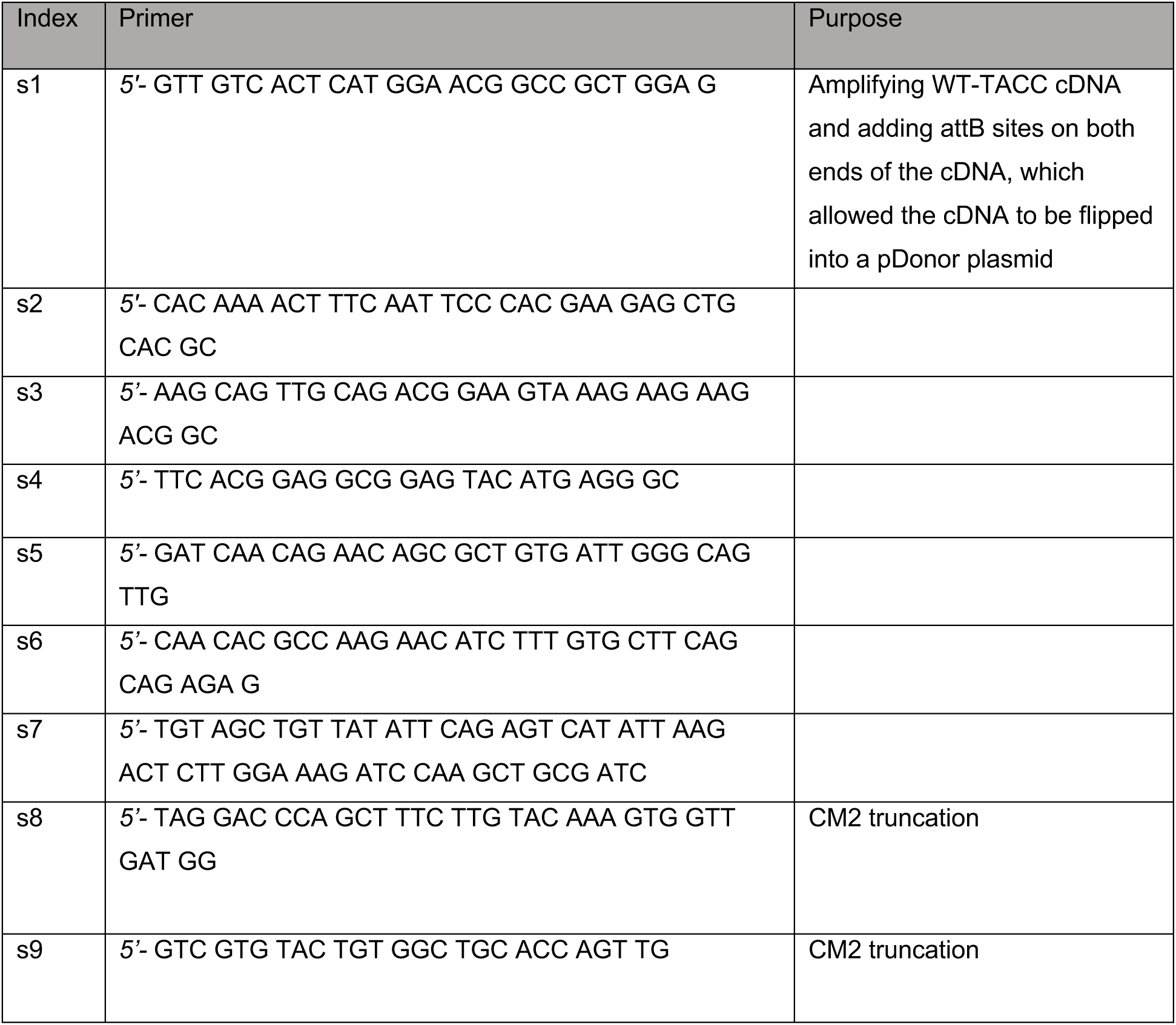
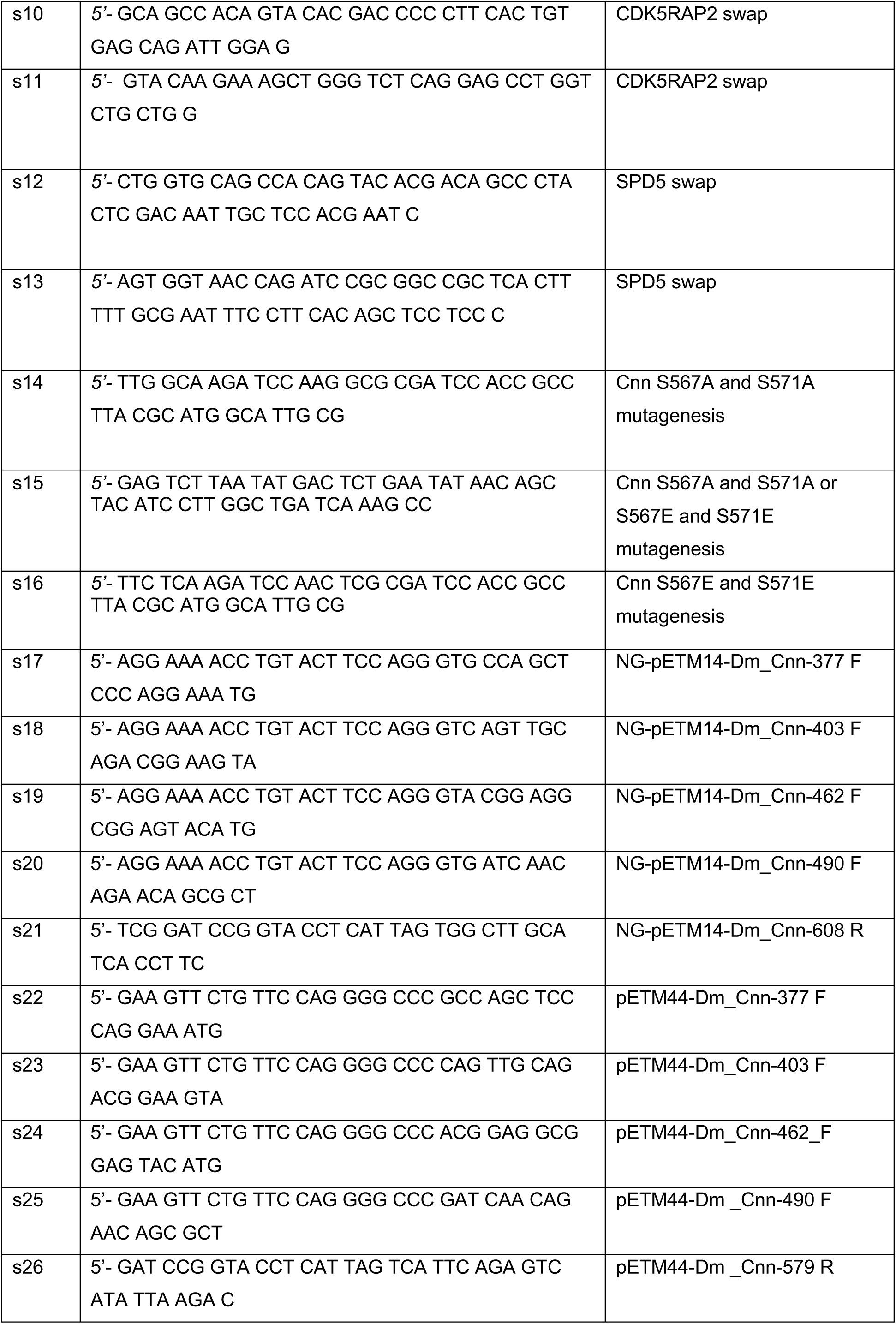
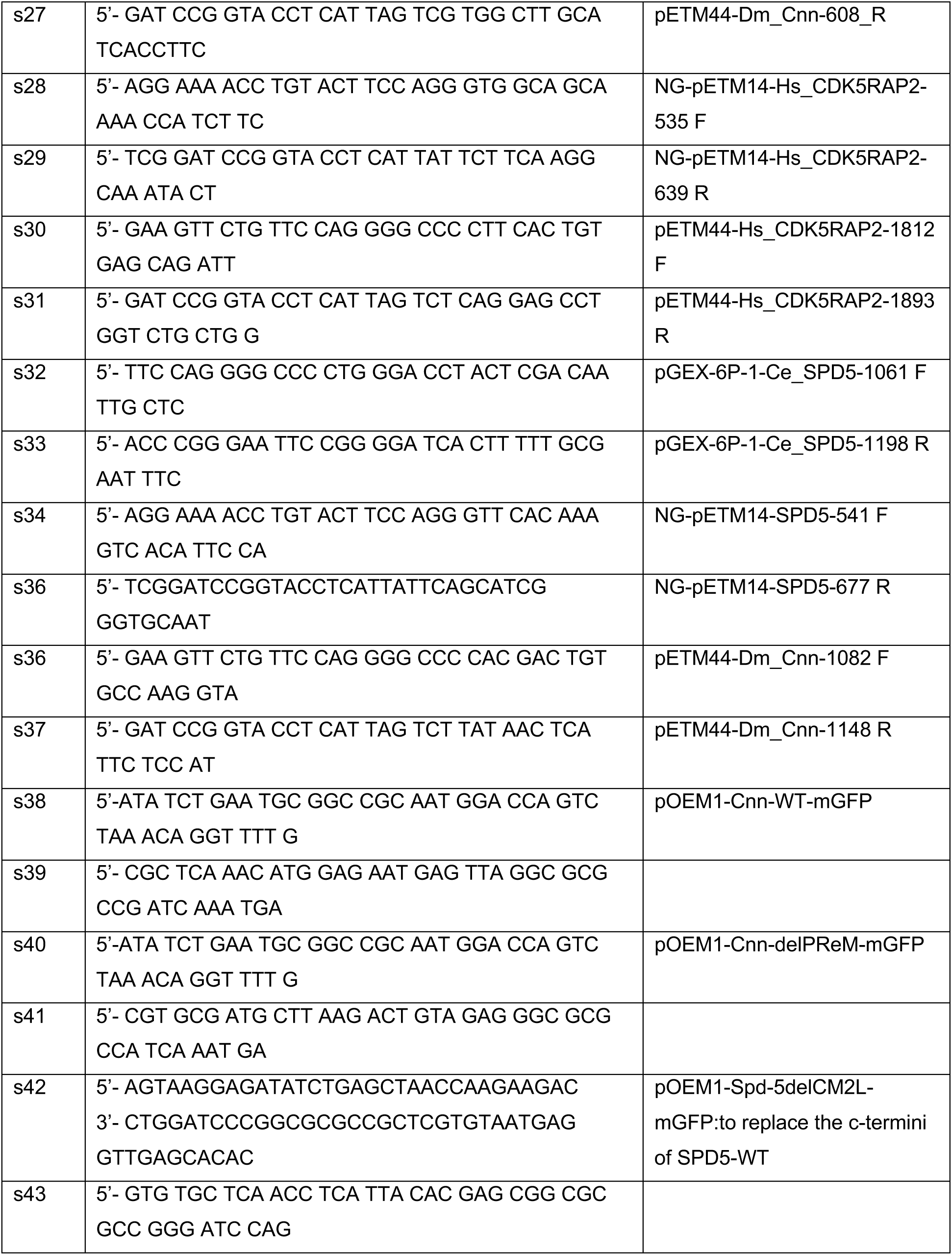
Primers used in this study.

**Table 4:** Plasmids used in this study.

| Index | Plasmid | Purpose | Source |
| --- | --- | --- | --- |
| p1 | pRNA-NG-Cnn | Plasmid contains non-phosphorylatable Cnn for RNA synthesis | Generated in this study |
| p2 | pRNA-NG-CnnΔ376-609 | | Generated in this study |
| p3 | pRNA-NG-CnnΔ376-402 | | Generated in this study |
| p4 | pRNA-NG-CnnΔ376-462 | | Generated in this study |
| p5 | pRNA-NG-CnnΔ376-490 | | Generated in this study |
| p6 | pRNA-NG-CnnΔ551-609 | | Generated in this study |
| p7 | pRNA-NG-CnnΔ583-609 | | Generated in this study |
| p8 | pRNA-NG-CnnΔCM2 | | Generated in this study |
| p9 | pRNA-NG-Cnn-[CDK5RAP2::CM2] |  | Generated in this study |
| p10 | pRNA-NG-Cnn-[SPD5::CM2] |  | Generated in this study |
| p11 | *pUbq-NG-Cnn-2A* | Plasmid encodes NeonGreen-tagged Cnn (S567A and S571A), under the control of a polyubiquitin promoter, for the generation of transgenic fly lines through *P*-element mediated transformation | Generated in this study |
| p12 | *pUbq-NG-Cnn-2E* | Plasmid encodes NeonGreen-tagged Cnn (S567E and S571E), under the control of a polyubiquitin promoter, for the generation of transgenic fly lines through *P*-element mediated transformation | Generated in this study |
| p13 | NG-pETM14-Dm_Cnn 377-608 |  | Generated in this study |
| p14 | NG-pETM14-Dm_Cnn 403-608 |  | Generated in this study |
| p15 | NG-pETM14-Dm_Cnn 462-608 |  | Generated in this study |
| p16 | NG-pETM14-Dm_Cnn<br>490-608 |  | Generated in<br>this study |
| p17 | pETM44-Dm_Cnn 377-608 |  | Generated in<br>this study |
| p18 | pETM44-Dm_Cnn 462-608 |  | Generated in<br>this study |
| p19 | pETM44-Dm_Cnn 490-608 |  | Generated in<br>this study |
| p20 | pETM44-Dm_Cnn<br>490-579 |  | Generated in<br>this study |
| p21 | pETM44-Dm_Cnn<br>490-579-2A | Plasmid encodes MBP -tagged Cnn 490-579<br>(S567A and S571A mutants) | Generated in<br>this study |
| p22 | NG-pETM14-Hs_CDK5RAP2- PReM<br>535-639 |  | Generated in<br>this study |
| p23 | pETM44-Hs_CDK5RAP2-CM2<br>1812 1893 |  | Generated in<br>this study |
| p24 | pGEX-6P-1-Ce_SPD5-CM2 1061-1198 |  | Generated in<br>this study |
| p25 | NG-pETM14-SPD5-PReM 541-677 |  | Generated in<br>this study |
| p26 | pETM44-Dm_Cnn-CM2<br>1082-1148 |  | Generated in<br>this study |
| p27 | pOEM1-MBP-long-linker_Cnn-WT-mGFP-long-linker-His10<br>(TH2036-pOCC502) |  | Generated in<br>this study |
| p28 | pOEM1-MBP-long-linker_Cnn-delPReM-mGFP-long-linker-His10 (TH2253-pOCC502) | Fragment-Cnn1-delPReM (403-608)-Bsu36I-PspOMI synthesised by Twist Bioscience in<br>2022 to generate plasmid encoding Cnn-delPReM construct | Generated in<br>this study |
| p29 | pOEM1-MBP-linker_Cnn-delCM2-mGFP-linker-His6<br>(TH1592-pOCC120) |  | Generated in<br>this study |
| p30 | pOEM1-MBP-linker_SPD-5-WT- mGFP-linker-His6<br>(TH0860-pOCC27) |  | (Woodruff *et al*,<br>2015) |
| p30 | pOEM1-MBP-linker_SPD-5-delPReM<br>(541-677)-mGFP-linker-His6 (TH2475-pOCC27) | Fragment PmII-AfIII (1. Spd5_del541-677_artSpel) with corresponding deletion<br>and artificial Spel (by silent mutagenesis)<br>was synthesised by GenScript Biotech in<br>2024 to generate the SPD-5delPReM<br>deletion construct. | Generated in<br>this study |
| p31 | pOEM1-MBP-linker_SPD-5-delCM2L(1161-1198)-<br>mGFP-linker-His6<br>(TH2077-pOCC27) |  | Generated in<br>this study |
| p32 | pOEM1-worm_PLK-1-mGFP-linker-His6<br>(TH1091-pOCC8) |  | (Woodruff *et al*,<br>2015) |
| p33 | pOEM1-MBP-long-linker_Dm_polo-T182D(CA)-mGFP-long-linker-His10<br>(TH2194-pOCC502) |  | Generated in<br>this study |
| p34 | pOEM1-MBP--<br>linker_Dm_polo-WT-mcherry-linker-His6<br>(TH1547-pOCC216) |  | Generated in<br>this study |
| p35 | pOEM1-MBP--<br>linker_Dm_polo-T182D(CA)-untagged-<br>linker-His6 (TH1553-pOCC-127) |  | Generated in<br>this study |

### Transgenic fly line generation

Transgenic fly lines were generated via random P-element insertion (injected, mapped, and balanced by ‘The University of Cambridge Department of Genetics Fly Facility’). For transgene selection, the *w^+^*gene marker was included in the transformation vectors and constructs were injected into the *w^1118^* genetic background.

### Molecular biology

Full-length *Drosophila* Cnn was cloned into the pRNA-mNeonGreen (pRNA-NG) destination vector by Gateway LR recombination (ThermoFisher Scientific) to generate pRNA-NG-Cnn (p1). Truncated variants were produced by linearising p1 with specific primer pairs and re-circularising using the KLD Enzyme Mix (NEB, M0554S). The following deletions were generated: residues 376–609 (p2), 376–402 (p3), 376– 462 (p4), 376–490 (p5), 551–609 (p6), and 583–609 (p7). A CM2 deletion mutant (p8) was obtained by linearising p1 with primers s8/s9. For domain-replacement constructs, the CM2 of Cnn was replaced with the CM2 of human CDK5RAP2 or the CM2-like region of C. elegans SPD-5, amplified with primers s10/s11 or s12/s13, respectively, and assembled into the s8/s9-linearised p8 using NEBuilder HiFi (NEB) to generate plasmids p9 and p10. Non-phosphorylatable (S567A/S571A, p11) and phosphomimetic (S567E/S571E, p12) Cnn mutants were created in pUbq-NG-Cnn by Q5 site-directed mutagenesis (NEB) using primers s14/s15 or s16/s15, respectively.

For bacterial expression, *Drosophila* Cnn fragments (residues 403–608, 377–608, 462–608 [WT or S567A/S571A], and 490–608) were cloned into a modified pETM14 vector encoding an N-terminal His₆–mNeonGreen tag. Equivalent constructs were generated for human CDK5RAP2 (residues 535–639) and C. elegans SPD-5 (residues 541–677). For MBP fusions, Cnn fragments (403–608, 377–608, 462–608, 490–608, and 490–579 [WT or S567A/S571A]) were cloned into NcoI-digested pETM44, encoding an N-terminal His₆–MBP tag, and the same design was used for CDK5RAP2 CM2 (residues 1812–1893). For GST fusions, C. elegans SPD-5 (residues 1061–1198) was cloned into BamHI-digested pGEX-6P-1, generating an N-terminal His₆–GST construct.

Full-length *Drosophila* Cnn, human CDK5RAP2, and *C. elegans* SPD-5 were synthesised (GenScript Biotech, Twist Bioscience, or Integrated DNA Technology) and subcloned into pOEM1-based baculovirus expression plasmids using NotI/AscI restriction sites. Primer sequences and plasmid details are provided in Tables 3 (primers s38-43) and 4 (plasmids p27-35).

### mRNA synthesis

For *in vitro* transcription, the constructs were digested and linearised by AscI (R0558S, NEB), and purified with PCR Purification Kit (QIAGEN). Around 1.6-3.2μg of digested and purified DNA was used to synthesise mRNA with the T3 mMESSAGE mMACHINE kit (AM1348, ThermoFisher Scientific). The mRNA product was purified with the RNeasy MinElute Cleanup Kit (74204, Qiagen). All RNAs were stored at −70°C. The final concentrations of RNA used in various experiments are described in the next section.

### Embryo collection and injection

Embryos were collected from plates (40% cranberry-raspberry juice, 2% sucrose, and 1.8% agar) supplemented with fresh yeast suspension. For live-imaging experiments, embryos were collected for 1h at 25°C, and aged at 25°C for 45–60 min. Embryos were dechorionated by hand, mounted on a strip of glue on either a 35-mm glass-bottom Petri dish with 14 mm micro-well (MatTek) and desiccated for 1 min at 25°C before covering with Voltalef oil (H10S PCTFE, Arkema) to avoid further desiccation.

### *In vitro* Condensate formation assay

Condensates of Cnn-WT, SPD-5-WT, and the corresponding deletion mutants were generated by diluting concentrated proteins, stored in high-salt buffer [50 mM HEPES pH 7.25, 500 mM KCl, 5% (v/v) glycerol, 1 mM DTT], into assay buffer of physiological ionic strength [50 mM HEPES pH 7.25, 150 mM KCl, 1 mM ATP/Mg²⁺, 2 mM TCEP] supplemented with 4% (v/v) polyethylene glycol (20 kDa). Prior to use, proteins were clarified by filtration through Ultrafree 0.22 µm PVDF filters (Merck Millipore) to remove residual aggregates. For imaging experiments, assay buffer was dispensed into 384-well ULA-coated microplates (Revvity PhenoPlate™, formerly CellCarrier Ultra), followed by the addition of the respective protein solutions to induce condensate formation.

### *In vitro* phosphorylation

Purified NG-SPD5-PReM was phosphorylated using *C. elegans* GFP-PLK1 in a time-course assay. Reactions (400 µl total volume) contained 100µM NG-SPD5-PReM, 10µl GFP-PLK1, 2 mM ATP, and 1× kinase buffer (from a 10× stock). Protein, PLK1, and ATP stocks were thawed on ice prior to use.

Reactions were incubated at 23 °C (optimal for C. elegans PLK1 activity) and sampled at T = 0, 1, 2, 3, and 4 h. At each time point, 10 µl was withdrawn for analysis by 20% SDS-PAGE, and ATP was replenished. Following the time course, samples were further purified on a pre-equilibrated Superdex 200 Increase 10/300 GL column (Cytiva) in buffer containing 20 mM Tris-HCl (pH 7.5), 150 mM NaCl, and 0.5 mM TCEP. Phosphorylation was confirmed by mass spectrometry (Advanced proteomics facility; Department of Biochemistry; University of Oxford).

### Live imaging and microscopy

Live embryo imaging was performed at 23 °C using a PerkinElmer ERS spinning disk confocal system mounted on a Zeiss Axiovet 200M microscope and controlled with Volocity software (PerkinElmer). Images were acquired with a 63×/1.4 NA oil objective (Carl Zeiss) using ImmersolT 518F immersion oil (RI = 1.518) to minimise spherical aberration. Fluorescence was detected with a 15-bit Orca ER CCD camera (Hamamatsu Photonics) operated at a gain of 200 V. Excitation was provided by 405, 488, 561, and 642 nm solid-state lasers (Oxxius S.A.). For dual-color acquisitions, red and green channels were imaged sequentially in each z-slice using the UltraVIEW ERS “Emission Discrimination” setting with 520 nm long-pass (green) and 620 nm long-pass (red) emission filters. Z-stacks were collected at 0.5 µm intervals, with imaging parameters (section number, time step, laser power, exposure) optimised for each experiment.

For estimation of protein saturation concentrations (C_sat_, Fig. 1D), imaging was performed at room temperature on a Nikon Eclipse Ti motorised stand equipped with a Yokogawa CSU-X1 spinning disk scan head and a Nikon Plan Apo 100×/1.45 NA oil objective. Samples were imaged with immersion oil (RI = 1.518) and illuminated using a 488 nm diode-pumped solid-state laser (Omicron) at 2% power. Emission was collected through a 525/30 band-pass filter. Images were acquired on an Andor iXON 897 EMCCD camera (16 µm pixel size) at 16-bit depth with no binning and a fixed amplifier gain of 200 V. Acquisition was controlled with NIS Elements software (v5.42.03), and imaging parameters were held constant across conditions for quantification.

Condensate diameters (Fig. 1F) were measured using an Olympus IXplore SpinSR microscope equipped with a Yokogawa CSU-W1 spinning disk and SoRa (Super-resolution via Optical Re-assignment) module. Samples were excited with a 488 nm diode-pumped solid-state laser at 50% intensity, and emission was collected with a 525/50 nm band-pass filter. Imaging was performed using a 3.2× SoRa magnification lens combined with an Olympus U ApoN O-TIRF 100×/1.49 NA oil objective, yielding a final system magnification of 320×. Fluorescence was recorded with a Hamamatsu ORCA-Fusion BT sCMOS camera (16-bit depth, 23.0 MHz pixel clock). For each condition, 20–22 randomly selected single-plane images were acquired per well. Image acquisition was managed with Olympus cellSens software.

For the experiment shown in Figure 3, purified NG-Cnn fragments (residues 403–608, 377–608, 462–608, or 490–608) were mixed with purified Cnn CM2 at a 1:10 molar ratio in a total reaction volume of 30 µl. In Figure 5, purified NG-Cnn 462–608 (WT or S567A/S571A mutant) was mixed with purified Cnn CM2 at the same ratio. As for Figure 6, NG-Cnn 462–608 was mixed with either Cnn CM2, CDK5RAP2 CM2, or SPD5 CM2. In Figure 7, NG-SPD5 541–677 (WT or *in vitro* PLK1-phosphorylated) was mixed with SPD5 CM2. CDK5RAP2 PReM (residues 535–639) was mixed with purified CDK5RAP2 CM2 (residues 1812–1893) for Figure 8.

In all reactions, red beads (FluoSpheres Carboxylate-Modified microspheres, 0.1 µm; Invitrogen F8801) were diluted 1:1000 in buffer and added to the reactions at a final dilution of 1:10,000. The reaction buffer contained 20 mM Tris-HCl (pH 7.5, room temperature), 150 mM NaCl, 0.5 mM TCEP, and 10 µM ZnCl₂. Reactions were incubated at room temperature for 30 min, after which 8 µl was applied to a slide (pipette tip cut to minimise shear) and immediately covered with an 18 × 18 mm coverslip. Networks were visualised using a Zeiss 880 microscope fitted with Airyscan detector using a 63x 1.4NA lens. Z-stacks were Airyscan-processed in 3D with a strength value of Auto (∼6) and visualised using a maximum intensity projection.

### *In vitro* half-FRAP (Fluorescence recovery after photobleaching) experiments

FRAP experiments on *in vitro* condensates were carried out using an Andor spinning-disk FRAPPA system mounted on an Olympus IX81 inverted motorised microscope equipped with a Yokogawa CSU-X1 scan head. Condensates were prepared from 3 µM protein (as in the droplet formation assay) and left to settle for 15 min before bleaching. Photobleaching was performed with a 488 nm laser line (via AOTF) at 15% power using the integrated FRAPPA module (30 ms dwell time, 2 repeats), targeting ∼50% of each condensate after acquiring 3–5 pre-bleach frames. Images were collected with an Olympus UPlanSApo 60×/1.2 NA water-immersion objective and recorded on an Andor iXon 897 EMCCD camera (SN 3880) at 30 ms exposure with EM gain set to 200 V. All acquisition and bleaching steps were controlled by Andor iQ software (v3.6).

### Image processing and analysis

For centrosome analysis, raw images were maximum-intensity projected, and background was corrected for uneven illumination using a custom Python script. Centrosomes were identified using the Crocker–Grier centroid-finding algorithm (Crocker & Grier, 1996) implemented in TrackPy (Allan et al., 2016). Signal-to-background thresholds were determined using Otsu’s method (Otsu, 1979). Centrosome area and integrated intensity were then calculated from the segmented regions.

For condensate quantification (Figure 1), single-plane fluorescence images were converted to grayscale, denoised with a Gaussian filter, and segmented by Otsu thresholding. Binary masks were generated, and small objects below a defined size cutoff were excluded. Masks were overlaid on the original images for quality control. Quantitative features—including condensate number, size distribution, mean fluorescence intensity inside versus outside condensates, and condensate fraction— were extracted and exported in CSV format for downstream analysis. Additional processing was performed in Fiji. Custom Python scripts are available on request.

For network quantification (Figures 3A-D; 5B; 6B; 8C), five grids of 100 z-stacks per coverslip were acquired using a 20×/0.8 NA Plan-Apochromat lens, with three coverslips analyzed per construct. Stacks were maximum-intensity projected and thresholded above background using a bespoke Python script, and the number of pixels above threshold was quantified per image.

Fluorescence recovery analysis followed the procedure described previously (Hubatsch *et al*, 2021) using custom software validated in Matlab 2023b or later. Briefly, droplet positions were registered in Fiji with the StackReg plugin, and recovery dynamics were extracted by drawing a three-pixel-wide stripe ROI perpendicular to the bleach boundary at each time point. The stripe was collapsed into 1D by averaging across its width and then used as the initial condition for FRAP analysis. A one-dimensional diffusion equation with experimentally determined boundary conditions was fit globally to the recovery curves of each droplet to obtain diffusion coefficients. To account for proteins that were only transiently rather than permanently bleached (Sinnecker *et al*, 2005), a uniform exponential recovery term was included in the model.

### Recombinant protein expression and purification

Recombinant proteins were expressed in Escherichia coli BL21 (DE3) cells cultured in LB broth. Protein expression was induced with 0.8-1 mM IPTG after cells reached an O.D of ∼0.6-0.7. Cells were left expressing overnight at 18°C before being harvested. Cells were lysed in a buffer containing 20mM TRIS, 500mM NaCl, 0.5mM TCEP using an Emulsiflex C5 homogeniser (Avestin) at 15,000 psi and at 4°C. The soluble protein fraction was then separated from the insoluble fraction by centrifugation at 5,000 rpm for 30 mins. Proteins of interest were then purified using Ni-NTA affinity chromatography (HisTrap HP; Cytiva), MBP affinity chromatography, GSTrap (Cytiva) depending on the tag, followed by size exclusion chromatography in buffer containing 20 mM Tris (pH 7.5), 150 mM NaCl, and supplied with 0.5 mM TCEP. The protein concentration was measured using a NanoDrop Spectrophotometer ND-1000 (ThermoFisher).

For SEC-MALS analysis and crystallisation trials or further use, the N-terminal tag was cleaved off using a lab purified GST-3C protease. The resulting untagged protein was further purified via reverse Ni-NTA chromatography and a second round of size exclusion chromatography in the same SE buffer.

### Full-length protein purification

The baculoviruses for protein expression in Sf9 cells were generated using the FlexiBAC system (Lemaitre *et al*, 2019), and subsequently 1%(vol/vol) of the respective P2 stocks of these baculoviruses were used to infect Sf+/Sf9 cultures for 72 h at 28 °C. The cells were harvested and resuspended in lysis buffer (50 mM HEPES pH 7.25, 500 mM KCl, 20 mM Imidazole, 5% Glycerol, 0.1% CHAPS, 1 mM MgCl_2_, 1 mM DTT, 30 µL Benzonase/100 mL Buffer, 1x protease inhibitor cocktail tablet EDTA-free/ 50 mL buffer), either flash-frozen in liquid N_2_, and stored at −80 °C till further purification, or directly used for purification. The cells were subjected to lysis on a LM-20 microfluidizer, with 2 passes on ice at 11000 p.s.i. The lysates were centrifuged in a Ti45 rotor in an XPN-90 (Beckman Coulter) ultra-centrifuge, at 29,000 rpm for 45 min at 4 °C. The supernatants were carefully decanted and filtered on a 0.22 µm PES filter and allowed to bind to 5 mL Protino Ni-NTA (Macherey-Nagel) columns, pre-equilibrated with the lysis buffer, at a rate of 2 ml/min. The columns were washed with 15 column volumes (CV) of lysis buffer, 5 CVs of 1 M KCl containing lysis buffer, and then proteins were eluted using 250 mM imidazole. The eluted solutions were then purified over either amylose resin (NEB) or 5 mL MBP-TRAP columns (Cytiva HP), pre-equilibrated with MBP-binding buffer (50 mM HEPES pH 7.25, 500 mM KCl, 5% Glycerol, 0.1% CHAPS, 1 mM DTT), followed by 5 CVs wash with the same buffer. Then 15 mM maltose containing buffer was used to elute the proteins from the MBP-TRAP columns. The obtained protein solutions were concentrated on 100 kDa amicon filters (Cytiva) and the N-terminal MBP and C-terminal His tags were cleaved off, using an in-house purified His-3C protease, before being subjected to a final size-exclusion step, either on a 24 mL Cytiva superose 6 increase 10/300 column or a 120 mL Cytiva HiLoad 16/600 Superose 6 column, pre-equilibrated with storage buffer (50 mM HEPES pH 7.25, 500 mM KCl, 5% Glycerol, 1 mM DTT). Peak fractions containing the proteins were pooled and further concentrated on 100 kDa amicon filters. The protein concentrations were determined at 280 nm and flash frozen for storage at −80 °C.

### Mass photometry

Mass photometry experiments were conducted using a Refeyn TwoMP instrument (Refeyn, Oxford, UK) to determine the molecular masses of full-length purified proteins. The technique relies on converting interferometric scattering contrast values into molecular weights using calibration curves generated from known protein standards, including bovine serum albumin (BSA) and human immunoglobulinG (IgG). Briefly, high-precision 24 × 50 mm coverslips were cleaned by alternating between isopropanol and Milli-Q water in a bath sonicator (3 minutes per wash), then dried under nitrogen. Silicone gaskets (Grace Biolabs, Cat. No. 103250) were cleaned in the same solvents (without sonication) and mounted centrally on the dried coverslips prior to loading onto the microscope stage. Standard stocks of BSA and IgG were diluted to 100 nM in MP dilution buffer (20 mM HEPES, 150 mM KCl, pH 7.25). For each measurement, 18 µL of this buffer was added to the gasket chamber and used to focus the microscope objective. Then, 2 µL of the 100 nM standard solution was added to achieve a final concentration of 10 nM. After gentle mixing, MP measurements were acquired immediately using AcquireMP software (Refeyn, v2.5.0). Contrast histograms were generated and fitted with Gaussian functions using DiscoverMP software (Refeyn, v2.5.0). The peaks corresponding to monomeric BSA (66 kDa), dimeric BSA (132 kDa), and IgG (150 kDa) were used to create a calibration curve for molecular weight determination. The same procedure was applied to the purified full-length protein samples. Molecular weight distributions were fitted with Gaussian curves using the previously generated calibration curve. A minimum of three technical replicates were performed for each protein sample to ensure reproducibility.

### Crystallisation, data collection and processing

#### Cnn 490-579 WT and 2A mutant

Crystallisation was performed using the sitting-drop vapor-diffusion method. Cnn 490-579 WT crystals were seen four days after setting the screening plates at 6mg/ml, 12mg/ml or 18mg/ml concentration in mother condition containing 0.8m potassium/sodium phosphate (pH 7.0). Tag cleaved Cnn 490-579 2A mutant purified protein was used to set crystallisation screens at a concentration of 10 mg/ml. Cnn 490-579 2A crystals grew in several conditions with the one giving the best diffraction growing in 1.32 M potassium/sodium phosphate pH 7 in the drop containing 75% protein sample and 25% of the buffer. Pipetting was carried out using mosquito LCP (ttplabtech). Crystal hits were screened by X-ray diffraction and data sets were collected at beamline I04 for Cnn 490-579 WT protein and I03 for the Cnn 490-579 2A mutant at the Diamond Light Source (DLS; Oxford, UK).

For Cnn 490-579 WT, Xia2 dials data reduction files was used for the initial structure model. As a homology model for phaser, a truncated version of CnnD490-K544(L535E) based on its complex structure with the Cnn C-terminus (PDB code: 5MW9) was used, which only retained the core dimer and lacked the termini (as well as its binding partner in the structure, the Cnn C-terminus). It was hypothesised that the dimeric core might be structurally conserved in CnnD490-N579. Indeed, a single solution with a translation function Z-score of 9.6 was obtained when using this search model. After some minor refinement involving changing a few side chain orientations, the model was fixed and phaser instructed to search for a straight, 20 amino acids long, polyalanine α-helix (adapted from PDB code: 5AL6). This resulted in 14 solutions, with the top hit having a translational function Z-score of 11.2. This cycle of placing a short stretch of a poly-alanine α-helix via phaser or molrep followed by refinement via Coot and refmac was repeated several times until no satisfactory placements were possible anymore. At this point, buccaneer was used with this partial solution and the expected protein sequence in order to build in the correct sequence of amino acids. The obtained model was used for refinement in phenix.refine including rigid body refinement to account for minor differences in the unit cell dimensions.

MTZ file from Cnn 490-579 2A mutant was processed with Xia2 3dii and were further indexed to space group P6_1_ using CCP4 suite. Structure was solved by molecular replacement using Phenix (Adams *et al*, 2010) using Cnn 490-579 WT structure (see above). Successful MR solutions were subjected to iterative manual building/refinement using Coot (Emsley & Cowtan, 2004) and Phenix.refine. Data processing and refinement statistics are summarised in Table 5.

**Table 5.** Data collection and refinment statistics of Cnn 490-579 WT and 2A mutant. Statistics from the highest resolution shell are indicated in parenthesis.

| Protein | Cnn<sub>490-579</sub> WT | Cnn<sub>490-579</sub> 2A |
| --- | --- | --- |
| Beamline | Diamond i04 | Diamond i03 |
| Wavelength | 0.9795 | 0.9763 |
| Resolution limit (Å) | 107.84 - 2.04 (2.07 - 2.04) | 46.45 - 2.00 (2.05 - 2.00) |
| Space group | P61 | P61 |
| Unit cell dimensions | 58.98, 58.98, 107.84 90, 90, 120 | 59.45, 59.45, 107.69 90, 90, 120 |
| Unique reflections | 13590 (634) | 14638 (1118) |
| Multiplicity | 11.2 (11.4) | 19.1 (12.5) |
| Completeness | 100.0 (96.8) | 100.0 (100) |
| I/sigma | 8.1 (1.4) | 15.5 (1.0) |
| Rmerge (%) | 15.8 (175.4) | 8.6 (226.2) |
| Rmeas (%) | 16.6 (183.7) | 8.8 (235.8) |
| Rpim (%) | 5.0 (54.3) | 2.1 (66.1) |
| CC1/2 | 1.0 (0.7) | 0.999 (0.499) |
| Processing pipeline | Xia2 dials (DLS) | Xia2 3dii (DLS) |

| Crystal | Cnn<sub>490-579</sub> WT | Cnn<sub>490-579</sub> 2A |
| --- | --- | --- |
| Resolution (Å) | 2.04 | 2 |
| Rwork/Rfree (%) | 20.55 / 21.82 | 22.9 / 27.2 |
| Asymmetric unit | Dimer | Dimer |
| Observed residues | Chain A: 493-576<br>Chain B: 492-577 | Chain A: 494-577<br>Chain B: 495-574 |
| % Data completeness (In resolution range) | 100 | 100 |
| Total number of atoms | 1452 | 1333 |
| Average B factors (Å) | 59 | 68 |
| Ramachandran statistics (%)<br>Favoured | 100 | 100 |
| Rotamer outliers (%) | 0 | 2.2 |
| PDB code |  | 9S7L |

### Protein structure prediction and visualisation

To identify the PReM region of *H. sapiens* CDK5RAP2, we first screened for potential interactions between the full N-terminus of CDK5RAP2 and its CM2 domain (residues 1812–1893) using AlphaFold3 with default settings (Abramson *et al*, 2024). Given prior evidence suggesting dimerisation, the input was designed to account for potential protein-protein interfaces. Although the initial prediction yielded a low ipTM score, the output was systematically analysed for plausible interaction interfaces.

Based on this analysis, the N-terminal region was fragmented into smaller domains while preserving its helical structural integrity. The ipTM score served as a preliminary filter for promising candidates. After iterative refinements of the input parameters, the CDK5RAP2 535–639 region was decided on as a potential binding target for the CM2 domain among others, despite its ipTM score remaining below 0.5. UCSF Chimera X-1.9 (Meng *et al*, 2023) was used for structural analysis and figure generation.

For the monomeric and dimeric structural predictions of the different Cnn deletions, CDK5RAP2 CM2 and SPD5 CM2, AlphaFold3 server was utilised with the default settings applied. UCSF Chimera X-1.9 was used for structural analysis and figure generation.

### Multiple Sequence Alignment (MSA)

Protein sequences of interest were obtained from UniProt (https://www.uniprot.org) using the following accession numbers: Cnn (*D.melanogaster*): UniProt ID P54623; CDK5RAP2 (*H.sapiens*): UniProt ID Q96SN8; and Spd5 (*C.elegans*): UniProt ID P91349.

MSAs were generated for the PReM region of Cnn across *Drosophila* species using MUSCLE with default parameters (gap opening penalty = −400, gap extension penalty = 0). The CM2 domain was aligned across a range of species using Clustal Omega with default settings (iterations = 3, guide tree clustering). All alignments were manually inspected for conserved motifs.

Sequence conservation was quantified using Jalview (v2.11.4.1) and visualised with colour mapping based on percentage identity (thresholds: ≥90% = dark blue, ≤30% = white).

### SEC-MALS analysis

SEC-MALS experiments were performed using a Superdex 75 10/300 Increase column or Superdex 200 10/300 increase column (Cytiva) and an AktaPure 25 System (Cytiva). The protein sample (100 μL) was loaded onto the gel filtration column and eluted with one column volume (24 mL) of buffer containing 20mM Tris pH 7.5, 150mM NaCl, and supplemented with 0.5mM TCEP, at a flow rate of 0.5 mL/min. The eluting protein was monitored using a DAWN HELEOS-II 18-angle light scattering detector (Wyatt Technologies), a U9-M UV/Vis detector (Cytiva), and an Optilab T-rEX refractive index monitor (Wyatt Technologies). Data were analysed by using Astra v7 (Wyatt Technologies) with a refractive increment value of 0.185 mL/g. Figures were generated using GraphPad Prism (version 10.4.2)

### Negative staining electron microscopy

Freshly glow-discharged TEM grids (C267, TAAB) were placed carbon-side down onto 10 µl droplets of sample containing the complex Cnn 490-608/Cnn CM21077-1148 at a 0.05mg/ml concentration resulting after SEC purification and incubated for 2 minutes. The grids (TAAB) were then blotted and incubated for 10 seconds on a 20 µl droplet of aqueous uranyl acetate 2% (w/v), then blotted and allowed to air dry. Grids were imaged using a JEOL Flash 120kV TEM equipped with a Gatan Rio camera.

### Molecular dynamics simulation

Simulations were run on the AlphaFold predicted structure of a PReM domain dimer (residues 490-608). This predicted structure was mutated in CHARMM-GUI to give structures for the 2A and 2E mutants (Jo *et al*, 2008). Charmm-GUI was also used to generate coarse grain systems for all constructs, using the Martini Solution Maker (Qi *et al*, 2015) with the martini3.0.0 force field. To match experimental conditions, amino acids protonation states were set to those at a pH of 7.5, and NaCl was added to the system at a concentration of 0.15M. Protein termini were set to neutral. Simulations were run in GROMACS (Bekker *et al*, 1993) using the default settings given by the CHARMM-GUI Martini Solution Maker, with two energy minimisation steps, one equilibration step, and a 200ns production run. Three simulations were run for each construct (2A, 2E and WT), with each equilibrated separately. Simulations were analysed by measuring distance between centre of gravity of the innermost (residues Arg 519-Lys 545) and second (residues Asn 559-Leu 586) helix for each chain over the course of each simulation.

### Statistical analysis

The details of statistical tests, sample size, and definition of the centre and dispersion are provided in individual Figure legends.

### Code and data availability

All raw image data will be deposited upon acceptance at The BioImage Archive (Hartley *et al*, 2022).

---

**Figure S1.**
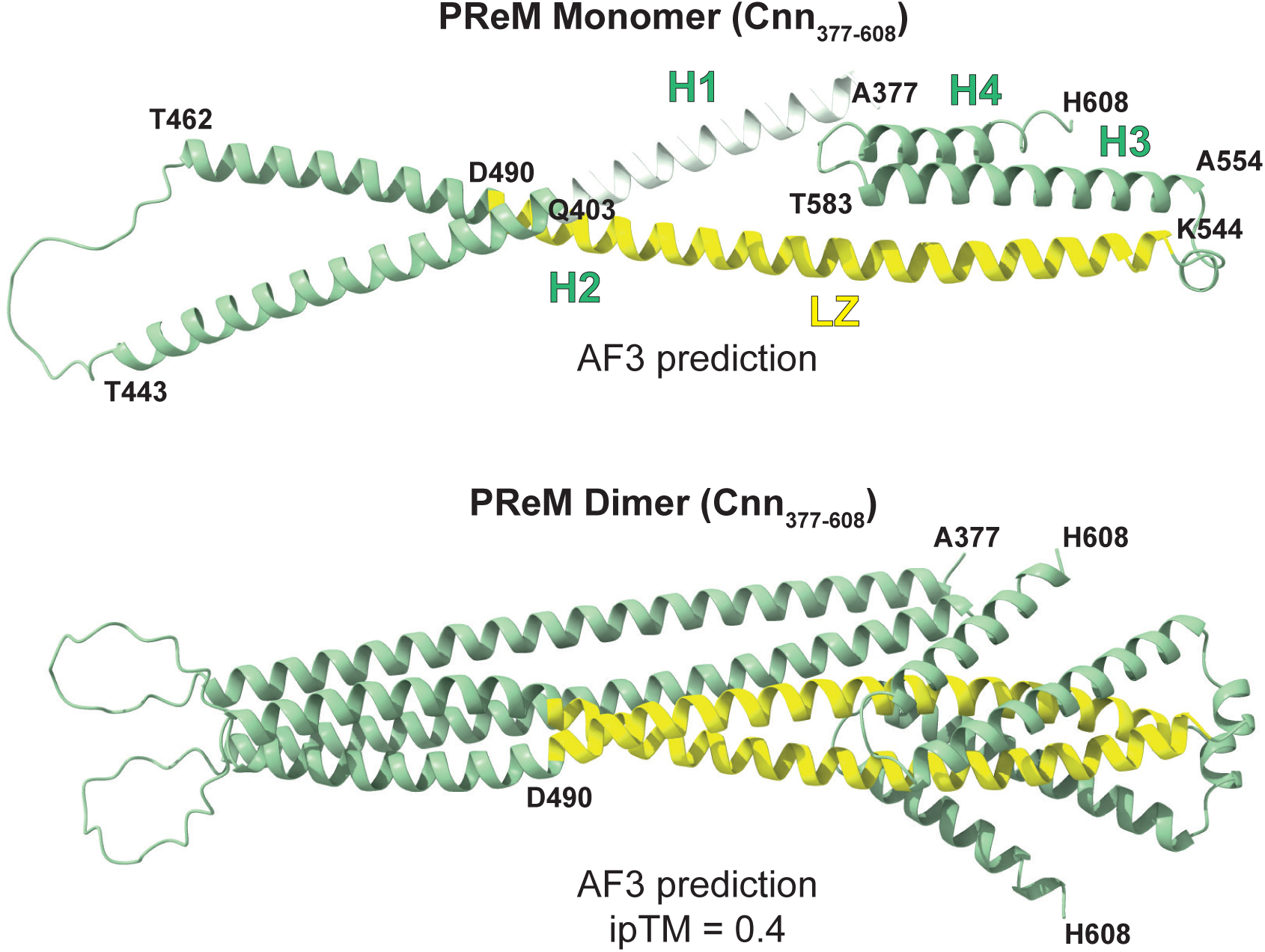
Predicted structures of the PReM domain. Ribbon representations of AlphaFold3 predictions of a PReM domain monomer (top) and dimer (bottom). Note that the PReM domain is almost certainly a dimer (see main text) but for simplicity we often show a schematic of a monomer to make the boundaries of various constructs easier to visualise.

**Figure S2.**
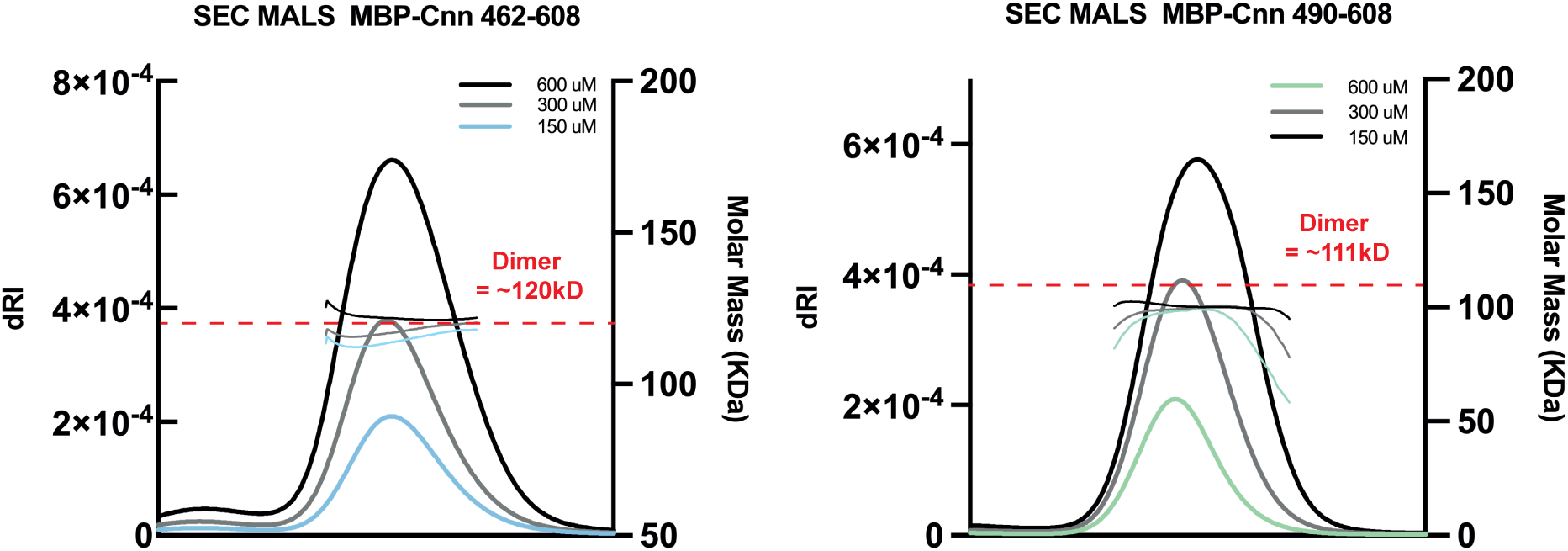
PReM domain constructs behave largely as dimers in solution. Traces of a SEC MALS experiment showing that MBP-fusions to two different PReM domain constructs (Cnn_462-608_ and Cnn_490-608_) behave largely as dimers in solution.

**Figure S3.**
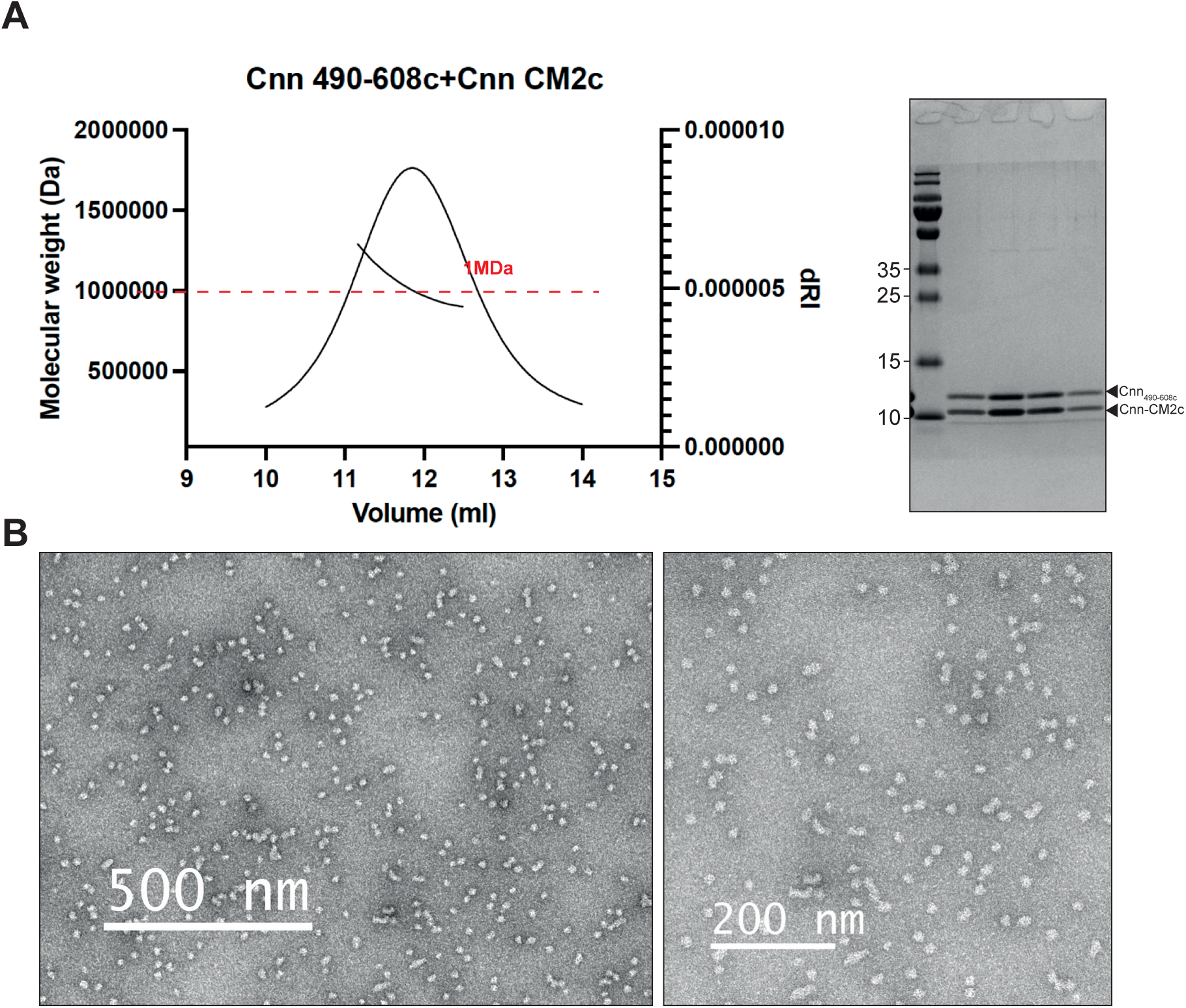
Cnn490-608 forms high MW complexes when mixed with CM2. **(A)** Traces of a SEC MALS experiment (and associated SDS gel) showing that Cnn_490-608_ forms high MW complexes of ∼ 1MDa when mixed with the CM2 domain *in vitro*. **(B)** Transmission EM images of the Cnn_490-608_::CM2 complexes.

**Figure S4.**
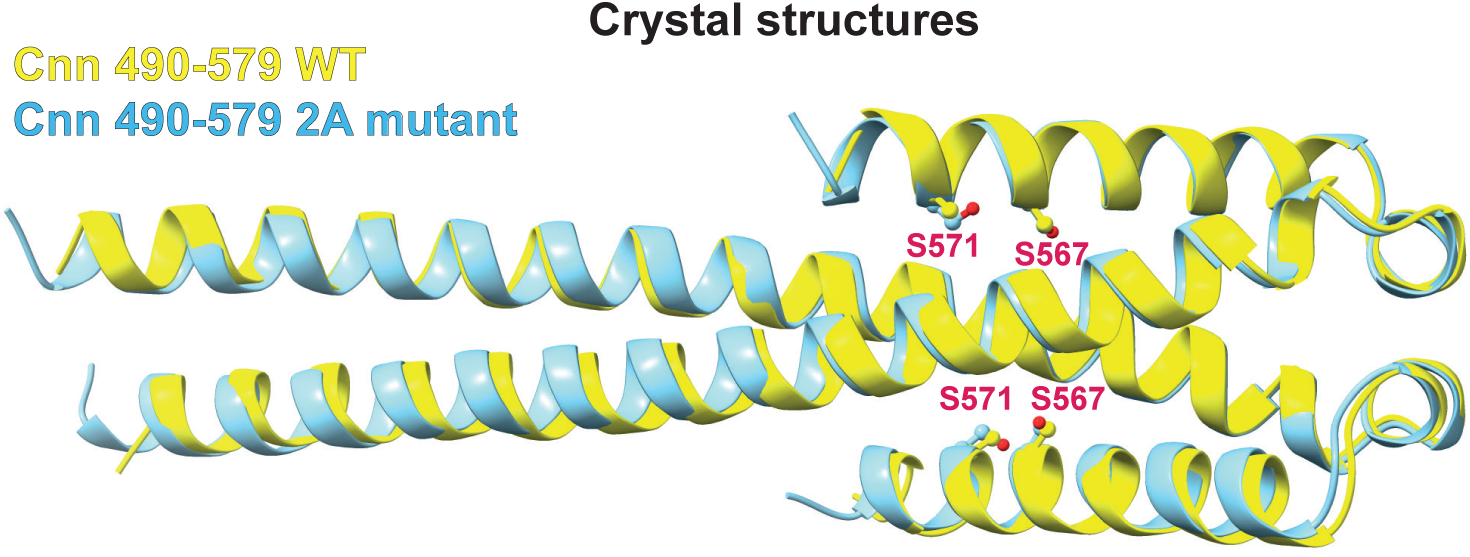
The Cnn-2A mutations do not perturb the helical hairpin structure of the PReM domain. An overlay of the crystal structures of the WT (*yellow*) and 2A mutant (*blue*) Cnn_490-579_ PReM domains, reveals that the mutations do not significantly alter the helical hairpin structure.

**Figure S5.**
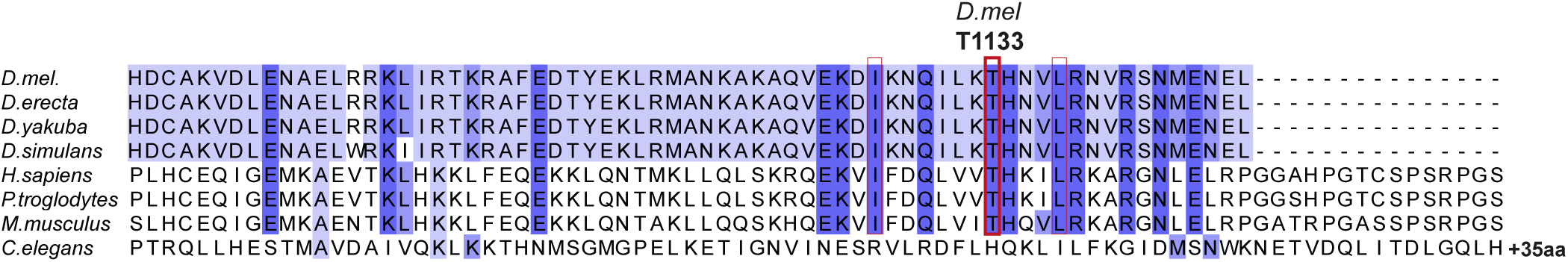
The *Drosophila* CM2 domain shows sequence homology to the CM2 domain of human CDKRAP2, but not to the putative CM2 domain of worm SPD-5. A multiple sequence alignment of the CM2 (and putative CM2) domains of Cnn, CDK5RAP2 or SPD-5 from various species. The conserved Thr (T1133 in *Drosophila*) is highlighted (*red* box); in *Drosophila*, T1133 binds to the PReM in the same region that is occluded by S567 in the PReM helical hairpin structure (Figure 4B), with S567 phosphorylation seeming to open the helical hairpin to allow CM2 to bind instead.

